# Phylogenetic divergence of GABA_B_ receptor signalling in neocortical networks over adult life

**DOI:** 10.1101/2024.06.04.597285

**Authors:** Max A. Wilson, Lewis W. Taylor, Soraya Meftah, Robert I. McGeachan, Tamara Modebadze, B. Ashan P. Jayasekera, Christopher J. A. Cowie, Fiona E. N. LeBeau, Imran Liaquat, Claire S. Durrant, Paul M. Brennan, Sam A. Booker

## Abstract

Cortical circuit activity is controlled by GABA-mediated inhibition in a spatiotemporally restricted manner. Much is known about fast GABA currents, GABA_B_ receptor (GABA_B_R) signalling exerts powerful slow inhibition that controls synaptic, dendritic and neuronal activity. However, little is known about how GABA_B_Rs contribute to circuit-level inhibition over the lifespan of rodents and humans. In this study, we quantitatively determine the functional contribution of GABA_B_R signalling to pre- and postsynaptic domains in rat and human cortical principal cells (PC). We find that postsynaptic GABA_B_R differentially control pyramidal cell activity within the cortical column as a function of age and species, and that these receptors contribute to co-ordination of local information processing in a layer- and species-dependent manner. These data directly increase our knowledge of translationally relevant local circuit dynamics, with direct impact on understanding the role of GABA_B_Rs in the treatment of seizure disorders.

**Highlights:**

- GABA_B_ receptor signalling displays age and species differences in cortex
- GABA_B_ receptor presynaptic inhibition is stronger in humans than rodents
- *In vitro* oscillations in human cortex are strongly regulated by GABA_B_Rs
- Levetiracetam enhances endogenous GABA_B_R signalling in human cortex

## Introduction

The transmission of information within cortical circuits requires integration of synaptic inputs to precisely control action potential output, which is true of all mammals. In mature neurons, this flow of excitatory information is largely orchestrated by the activation of GABA receptors which reduce neuronal activity by providing hyperpolarising or shunting inhibition to neuronal membranes in a cell-compartment specific manner (Mody and Pearce, 2004). While much is known about GABA_A_Rs, determining how GABA_B_Rs contribute to neuronal function across brain development is critical, as they are implicated in the aetiology of seizures and epilepsy in humans (Princivalle et al., 2003; Sheilabi et al., 2018; Teichgräber et al., 2009) and preclinical rodent models (Codadu et al., 2019; Schuler et al., 2001; Straessle et al., 2003). The mammalian neocortex displays both a columnar and laminar structure, with 6 defined layers extending from superficial layer 1 (L1) to deep L6. Layers 2-6 are populated with principal cells (PCs) which possess dendrites that terminate locally and in L1 (DeFelipe et al., 2002). While cortical columns and lamina organisation are generally common to all mammals, increased human cortical thickness allows greater subdivision of labour and expansion of cell types, not observed in rodent neocortex (Berg et al., 2021). Beyond PCs, the neocortex contains a diverse population of inhibitory interneurons (INs) which make up 44% of neurons in humans (DeFelipe et al., 2007); the types of which are broadly conserved with mice (Bakken et al., 2021). Local INs form dense connections with PCs and each other to produce strong inhibition of their target cellular compartments via monosynaptic activation of GABA_A_ receptors (GABA_A_Rs; (Campagnola et al., 2022)), and metabotropic GABA_B_Rs (Kohl and Paulsen, 2010; Kulik et al., 2018).

GABA binds with high affinity to GABA_B_Rs following heterosynaptic spill-over from nearby inhibitory synapses (Booker et al., 2013; Scanziani, 2000; Urban-Ciecko et al., 2015), which activates inwardly-rectifying K^+^ (Kir3) channels to hyperpolarise neuronal membranes (Bettler et al., 2004). This K^+^-conductance hyperpolarises dendrites over a long temporal window, but also displays prominent attenuation in electrotonically large neurons (Degro et al., 2015; Eder et al., 2001; Schulz et al., 2021). Somatodendritic GABA_B_R activation directly reduces spike output (Ault and Nadler 1983), dendritic integration (Palmer et al., 2013; Pérez-Garci et al., 2013), and network oscillations (Booker et al., 2020; Brown et al., 2007). In presynaptic compartments, GABA_B_Rs inhibit voltage-gated Ca^2+^-channels (VGCCs; (Bettler *et al*., 2004)) to regulate neurotransmitter release (Kanigowski et al., 2023; Kulik *et al*., 2018). However, GABA_B_Rs are not uniformly expressed in rodent brains, with a gradient of expression in both neocortex and hippocampus (Degro *et al*., 2015; Fritschy et al., 1999; Murer et al., 1997), a pattern which is not observed in post-mortem human brain tissue (Berthele et al., 2001; Billinton et al., 2000), despite the presence of functional GABA_B_Rs (Deisz, 1999; Teichgräber *et al*., 2009). A major confound to human studies is whether tissue was collected from seizure-free individuals, as complex GABA_B_R expression patterns are observed in different epilepsy patient groups (Princivalle *et al*., 2003; Sheilabi *et al*., 2018; Teichgräber *et al*., 2009). Regardless, the role of GABA_B_Rs in controlling cortical neuron and circuit function has not been fully characterised across the adult lifespan. We hypothesise that GABA_B_R signalling in the human neocortex may display similarity to that of rodents in an age dependent manner, and be affected by a clinical history of seizures and/or anti-seizure medications.

To address this, we directly quantified functional GABA_B_R signalling in the neocortex from rats and humans using whole-cell patch-clamp and extracellular local field potential (LFP) recordings. We show that while somatic GABA_B_R signalling is low in L5 PCs in juvenile rats, this increases into adulthood. Adult human cortical neurons display distinct GABA_B_R current densities and possess stronger presynaptic inhibition which combines to attenuate neuronal activity in the whole cortical column. At the circuit level, GABA_B_Rs control the strength and synchrony of neuronal oscillations in human cortex. Finally, we show that GABA_B_R signalling is enhanced in patients who receive the anti-seizure medication levetiracetam – irrespective of whether they experienced seizures. These findings highlight important phylogenetic differences in inhibitory signalling between rodents and humans, with direct implication for targeting GABA_B_Rs in the treatment of seizure disorders.

## Results

### Divergent baseline excitability of seizure-free humans and rat neocortex

Human neurons are known to possess some divergence in electrophysiological properties from rodents (Eyal et al., 2016; Kalmbach et al., 2018). To confirm this, we first compared the basal physiology of L2/3 and L5 neurons from primary somatosensory (S1) cortex of 1- and 6+ month old rats to human cortical neurons (**Figure 1A-C**). Human data was obtained from recordings from 49 individuals who were aged between 28 and 77 years old (Median: 60 years). All human brain tissue was obtained as ‘access tissue’ from individuals undergoing tumour resection. As seizure or anti-seizure medication history may influence neuronal excitability and neurotransmitter signalling, we first analysed data from only those patients that were seizure free and had not received anti-seizure medications (N=26 cases; **Supplementary Table 1**).

**Figure 1:**
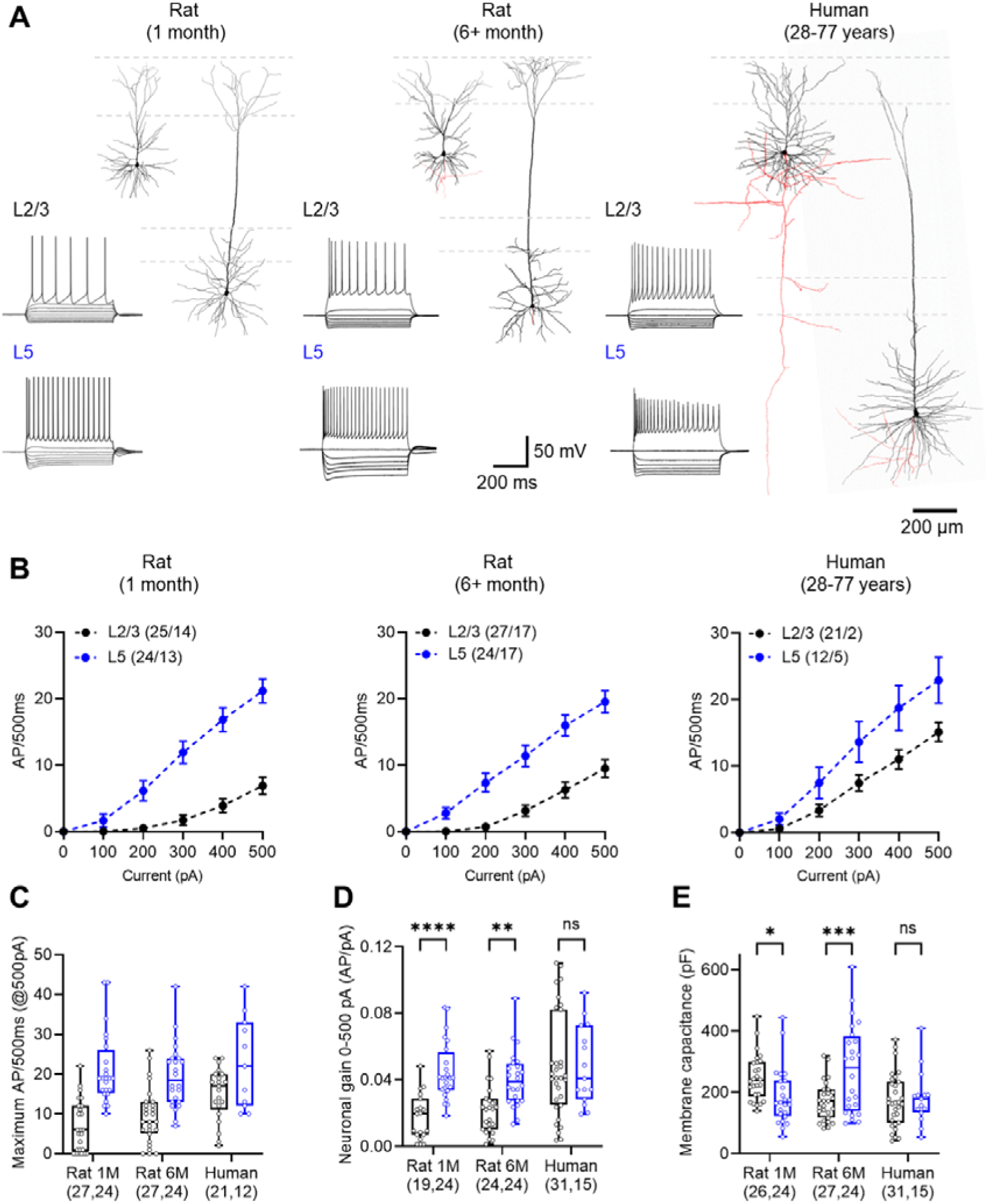
Age and species dependent changes in neuronal excitability. (**A**) Example reconstructions of L2/3 and L5 neurons from rat and human neocortex. Somatodendritic axis (black) and axons (where present, red) are shown with respect to cortical layering (grey dashed lines). Inset, voltage response of example cells in response to -500 to +500 pA current injections (100 pA steps, 500 ms duration). (**B**) Current voltage responses of L2/3 (black) and L5 (blue) neurons from 1-month (left) and 6+ month rats (middle) and human neocortex (right). (**C**) Peak AP discharge from recorded neurons between layers (F_(1, 116)_ = 50.14; P<0.0001; 2-way ANOVA [layer]), age and species (F_(2, 116)_ = 3.455, P=0.035; 2-way ANOVA [age+species]). The number of cells recorded per cell type are shown underneath (L2/3, L5). (**D**) Neuronal gain plotted for recorded neurons, which displayed layer and species/age differences (F_(2, 130)_ = 4.462, P=0.013; 2-way ANOVA [age+species x layer]). (**E**) Membrane capacitance of recorded neurons, which displayed layer and species/age differences (F_(2, 130)_ = 9.737, P=0.0001; 2-way ANOVA [age+species x layer]). All data is shown as either mean ± SEM (B) or 25 - 75% boxplot with median indicated ± range (C, D, E). All data are overlain by responses of individual cells. Statistics shown from 2-way ANOVA, with post-tests shown as: ns – p>0.05, * - P<0.05, ** - P<0.01; *** - P<0.001, **** - P<0.0001 from Holm-Sidak post-tests.

L2/3 neurons typically displayed fewer action potentials (APs) than L5 counterparts in response to depolarising current steps, irrespective of being from rat or human brain tissue (**Figure 1B**). However, when comparing either the peak AP discharge (**Figure 1C**) or neuronal gain (**Figure 1D**), L2/3 PCs displayed increased excitability in a species- and age-dependent manner, which was not observed in L5 PCs. This increase in L2/3 activity was unlikely due to changes in cell size, as although membrane capacitance did show differences over age, L2/3 PCs of 6+ month old rats were largely similar to humans; meanwhile L5 PCs showed divergence between these groups (**Figure 1E**). A summary of human intrinsic physiology is shown in **Supplementary Table 2**. As many of the key electrophysiological properties of L2/3 neurons show age-dependent changes in rats, we asked if such changes are present during adulthood in humans. To address this, we compared the age of patients to the electrophysiological properties of individual neurons. We found no significant correlation of any tested neuronal parameter with age **(Supplementary** Figure 1).

### Divergent functional GABA_B_R-mediated currents in human and rat neocortex

The human cortex is known to express functional GABA_B_Rs (Deisz, 1999) that may show age dependent changes in expression at total protein levels (Pandya et al., 2019). Whether the function of these receptors deviate from those of rodents remains largely unexplored. To determine the functional expression of GABA_B_Rs in neurons, we performed whole-cell recordings from identified L2/3 and L5 PCs from rats and humans in the presence of blockers of AMPA, NMDA, and GABA_A_ receptors leaving GABA_B_R signalling intact. Following baseline recording we applied the selective GABA_B_R agonist baclofen to the bath to activate all receptors on recorded neurons, followed by the potent and selective GABA_B_R antagonist CGP-55,845 (CGP, 5 μM, 5 minutes).

In 1 month-old rats, we observed a large outward current in L2/3 PCs following baclofen application, which was blocked by subsequent application of 5 µM CGP (Figure 2A). As expected from previous reports (Palmer *et al*., 2013; Schulz *et al*., 2021), we observed 59% lower whole-cell currents in L5 PCs in 1-month old rats when quantified as the peak baclofen current (t_(10,11)_=5.3, p=0.002, Holm-Sidak test; Figure 2B). In 6+ month-old rats, L5 neurons had total currents largely similar to L2/3 (t_(20,16)_=0.97, p=0.336, Holm-Sidak test). In human neurons, we likewise observed minimal difference between L2/3 and L5 neurons (t_(28,10)_=0.73, p=0.469, Holm-Sidak test). As cell capacitance (a proxy for electrotonic size) differed between cell types (Figure 1E), we next normalised baclofen whole-cell currents to measure capacitance for each cell (Figure 2C). We found in 1 month-old rats, the GABA_B_R current density was 63% lower in L5 PCs compared to L2/3 (t_(10,11)_=5.3, p<0.0001, Holm-Sidak test). In 6+ month old rats, L5 PCs had slightly lower (−26%) baclofen current-density, which was not significantly different from L2/3 PCs (_(20,16)_=1.8, p=0.073, Holm-Sidak test). Indeed, this effect appeared due to an age dependent decline in GABA_B_R signalling, specific to L2/3 cells, from early adolescence into adulthood in rats (F_(2, 88)_ = 5.743, *p*=0.0045, 2-way ANOVA [age+species]). In human cortical neurons, we found that L5 PCs had slightly (+18%) higher baclofen current density, than L2/3 PCs (t_(28,10)_=0.69, p=0.49, Holm-Sidak test). To test whether GABA_B_R functional currents are impacted by age in human neurons, as has been suggested in proteomic studies (Pandya *et al*., 2019), we compared the effect of age on GABA_B_R current-density over human adulthood. In contrast to rats, we observed no age-dependent decline in GABA_B_R current density between the ages of 28 and 77 for either L2/3 PCs or L5 PCs (Figure 2D).

**Figure 2:**
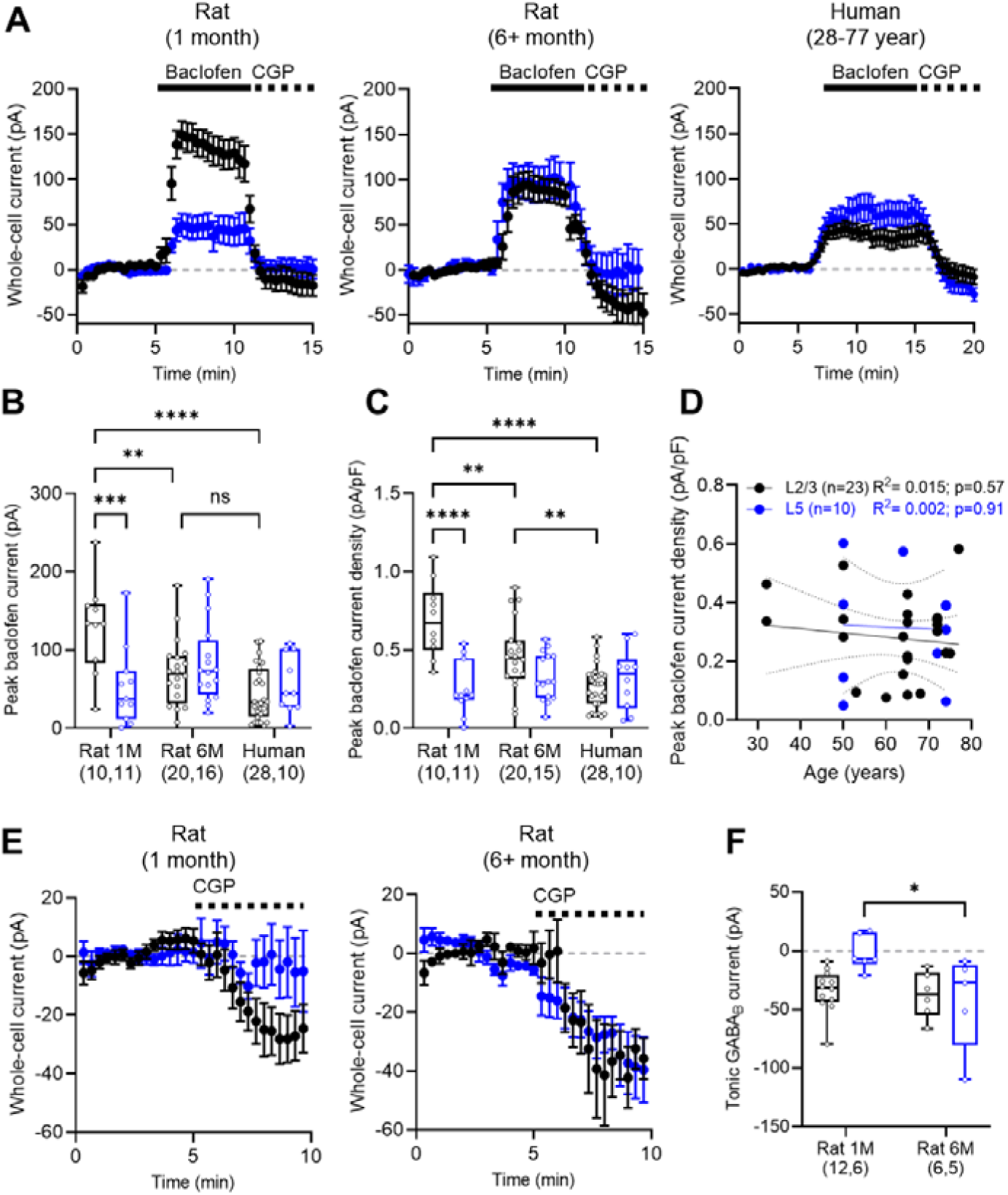
Age and cell-type specific differences in postsynaptic GABA_B_R currents in rats and humans. (**A**) Time-course plots from L2/3 (black) and L5 (blue) PCs from 1-month (left) and 6+ month (middle) rats, and human (right) neocortex, following bath-application of 10 μM baclofen (solid line) and 5 μM CGP (dashed line). The baseline current level is shown for reference (grey dashed line). (**B**) Peak baclofen current measured immediately following bath wash-in from recorded neurons (F_(2, 89)_ = 7.84, P=0.0007; 2-way ANOVA [age+species x layer]). (**C**) Peak baclofen current density from recorded neurons (F_(2, 88)_ = 10.21, P=0.0001; 2-way ANOVA [age+species x layer]). (**D**) Scatter plot of baclofen current-density from human L2/3 (black) and L5 (blue) neocortical neurons plotted against age (years). R^2^ values from linear regression are shown on graphs. (**E**) Time-course plots from L2/3 (black) and L5 (blue) PCs from 1-month (left) and 6+ month (right) rats following bath-application of 5 μM CGP (dashed line). (**F**) Tonic GABA_B_R mediated currents from rat neurons (F_(1, 25)_ = 4.220, P=0.05; 2-way ANOVA [age+species x layer]). All data is shown as either mean ± SEM (A, E) or 25 - 75% boxplot with median indicated ± range (B, C, F). All data are overlain by responses of individual cells. Statistics shown from 2-way ANOVA, with post-tests shown as: ns – p>0.05, * - P<0.05, ** - P<0.01; *** - P<0.001, **** - P<0.0001 from Sidak’s post-tests.

In some recordings, cells displayed negative whole-cell currents relative to baseline when CGP was bath applied (Figure 2A); indicative of a tonic GABA_B_R current. To directly test this, in a subset of recordings we applied CGP alone to rat slices, without prior baclofen application (Figure 2E). Consistent with baclofen whole-cell currents, we observed larger tonic currents in L2/3 compared to L5 PCs in 1 month-old rats, which was absent in 6+ month-old rats (Figure 2F; F_(1, 25)_ = 4.22, *p*=0.05, 2-way ANOVA [age x layer]), supporting the notion that layer-wise differences in GABA_B_R signalling are normalised by adulthood. These data reveal that functional GABA_B_Rs undergo an early life down-regulation in rats, with the greatest currents observed at younger ages. In seizure-free adult humans GABA_B_R mediated currents were similar between L2/3 and L5 PCs at all ages examined.

We next determined whether GABA_B_R-mediated currents in L2/3 and L5 neurons were driven by endogenous GABA release. To achieve this, we delivered short trains of high-frequency stimulation to the L1/2 border (5 stimuli, 200 Hz, 50 V) in the presence of the same ionotropic receptor blockers as above. In L2/3 neurons from all species and ages, electrical stimulation elicited large amplitude, slow IPSCs, which were sensitive to application of CGP-55,845 (Figure 3A). IPSCs elicited in 1 month-old rats agreed with baclofen data (Figure 2), with typically larger GABA_B_R-mediated slow-IPSCs in L2/3 PCs compared to L5 PCs. This was not the case for 6+ month rats and human cortical neurons, where slow-IPSCs were similar in amplitude in L2/3 and L5 PCs (Figure 3B). The combination of GABA_B_Rs, effector channels, and auxiliary proteins can alter receptor kinetics (Fritzius et al., 2017), as well as electrotonic properties (Degro *et al*., 2015), thus we next measured slow-IPSC kinetics (Figure 3C). We measured the 20-80% rise time of IPSCs, which were 51% faster in L5 PC than L2/3 PCs in 1 month old rats (t_(18,5)_=2.2, p=0.03, Holm-Sidak test). Such an effect was absent in 6+ month old rats (t_(9,5)_=1.4, p=0.16, Holm-Sidak test) and reversed in the human cortex, with the latter showing 65% longer 20-80% rise-time for L5 PCs compared to L2/3 (t_(10,4)_=4.0, p=0.0003, Holm-Sidak test). The decay time-constants of evoked slow-IPSCs were more variable, particularly in adult rats and humans, with a tendency to longer decay times in L5 neurons (Figure 3D).

**Figure 3:**
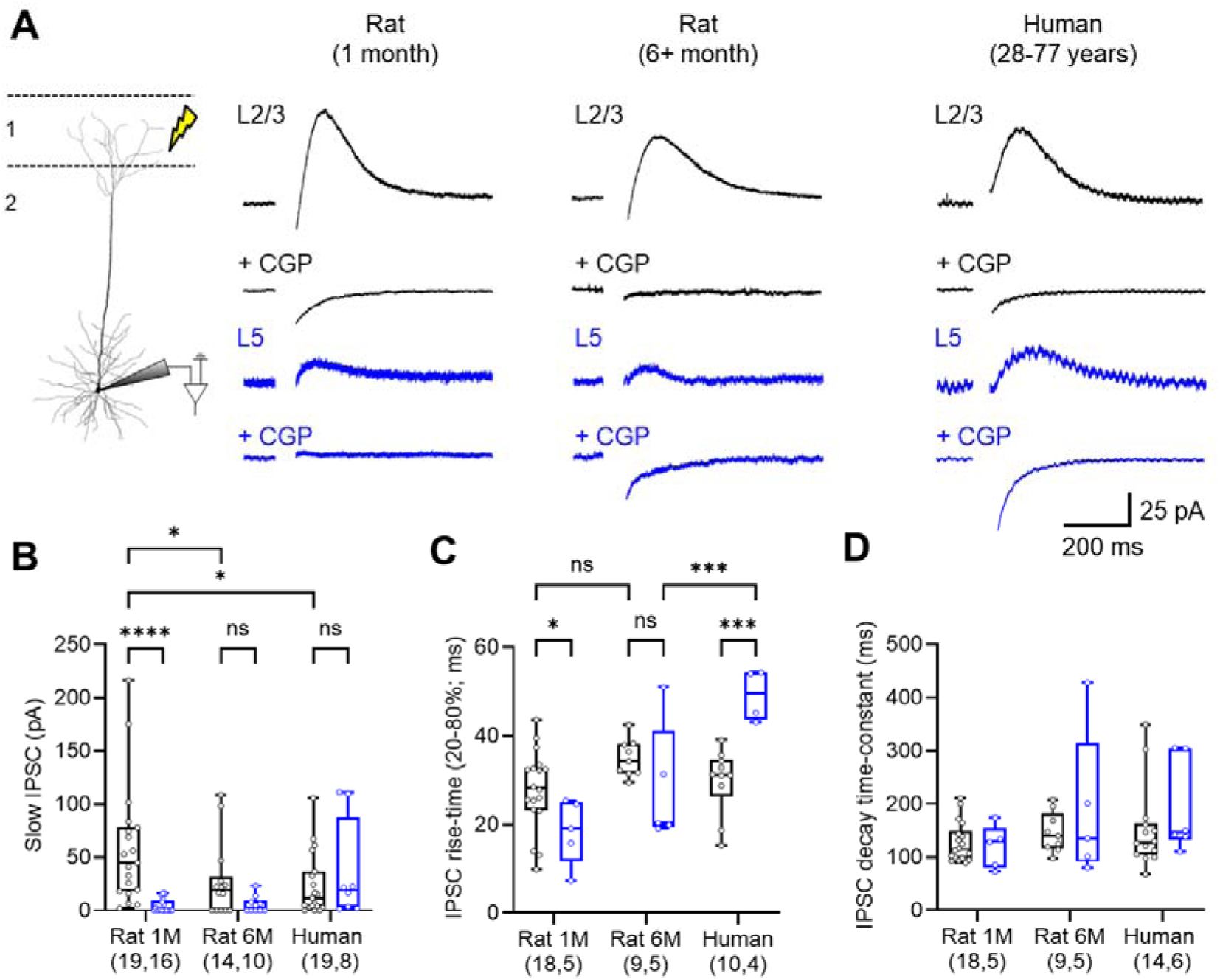
GABA_B_Rs are activated by endogenous GABA release with age and species-specific kinetic differences. (**A**) Left, Schematic of recording set up for identification of GABA_B_R-mediated IPSCs in neocortical PCs. Evoked IPSCs recorded in the presence of AMPA, NMDA, and GABA_A_ receptor antagonists/blockers (upper) and confirmed as GABA_B_R-mediated following bath application of 5 μM CGP (lower) from L2/3 (black) and L5 (blue) PCs from 1- and 6+ month old rats, and humans. (**B**) Quantification of IPSC amplitude in all groups, which differed by layer and age/species (F_(2, 74)_ = 4.00, P=0.02; 2-way ANOVA [age+species x layer]). (**C**) 20-80% rise time of pharmacologically isolated GABA_B_R IPSCs in neocortical neurons (F_(2, 44)_ = 5.37, P=0.008; 2-way ANOVA [age/species]). (**D**) Measured decay time-constants of GABA_B_R IPSCs did not differ across age or species (F_(2, 44)_ = 2.04, P=0.14; 2-way ANOVA [age+species]) or layer cell-type (F_(1, 44)_ = 0.45, P=0.51; 2-way ANOVA [layer]). All data is shown as 25-75% boxplots with median indicated ± range. All data are overlain by responses of individual cells. Statistics shown from 2-way ANOVA, with post-tests shown as: ns – p>0.05, * - P<0.05, ** - P<0.01; *** - P<0.001, **** - P<0.0001 from Sidak’s post-tests.

These data show that GABA_B_Rs are reliably recruited by GABA release in human and rat cortical circuits, giving rise to slow-IPSCs which display species differences in size and onset kinetics.

### GABA_B_Rs display developmentally divergent contributions to perisomatic inhibition

GABA_B_Rs in juvenile cortical neurons have been shown to largely localise to distal dendritic compartments, thus may have minimal direct influence on perisomatic excitability (Palmer *et al*., 2013; Schulz *et al*., 2021). Given age and species dependent differences in observable GABA_B_R-mediated currents, we asked if these receptors differentially modulate somatic excitability. For this we focally applied baclofen (50 μM, 200 ms) directly to the perisomatic region of neurons (Figure 4A). Perisomatic baclofen puff application gave rise to outward currents in recorded L2/3 and L5 cells in rats and human neurons (Figure 4B), which were blocked by bath application of CGP (Figure 4D**, 4F, 4H**). To determine whether perisomatic GABA_B_Rs were able to control neuronal activity we puff applied baclofen to neurons entrained to tonic firing (rheobase + 25 pA). Focal baclofen application to the perisomatic region consistently reduced the action potential discharge of L2/3 cells, independent of age or species. The observed reduction in AP discharge was blocked by CGP application (Figure 4C**, 4E, 4G**). The same focal GABA_B_R activation in L5 pyramidal cells inhibited discharge in 1-month old rats, less so than for L2/3 PCs (p=0.002, Mann-Whitney test; Figure 4D). In 6+ month old rats, the same baclofen puff produced near complete loss of firing in L5 PCs, comparable to L2/3 PCs (p=0.65, Mann-Whitney test; Figure 4F); which was also observed in humans (p=0.87, Mann-Whitney test; Figure 4H). These data confirm that GABA_B_Rs located in the perisomatic compartment more efficiently control the output of L5 neurons in adult human and rat cortex. These age dependent effects are likely due to increased peri-somatic localisation of the receptor and the hyperpolarisation produced by its activation in L5 PCs over later life.

**Figure 4:**
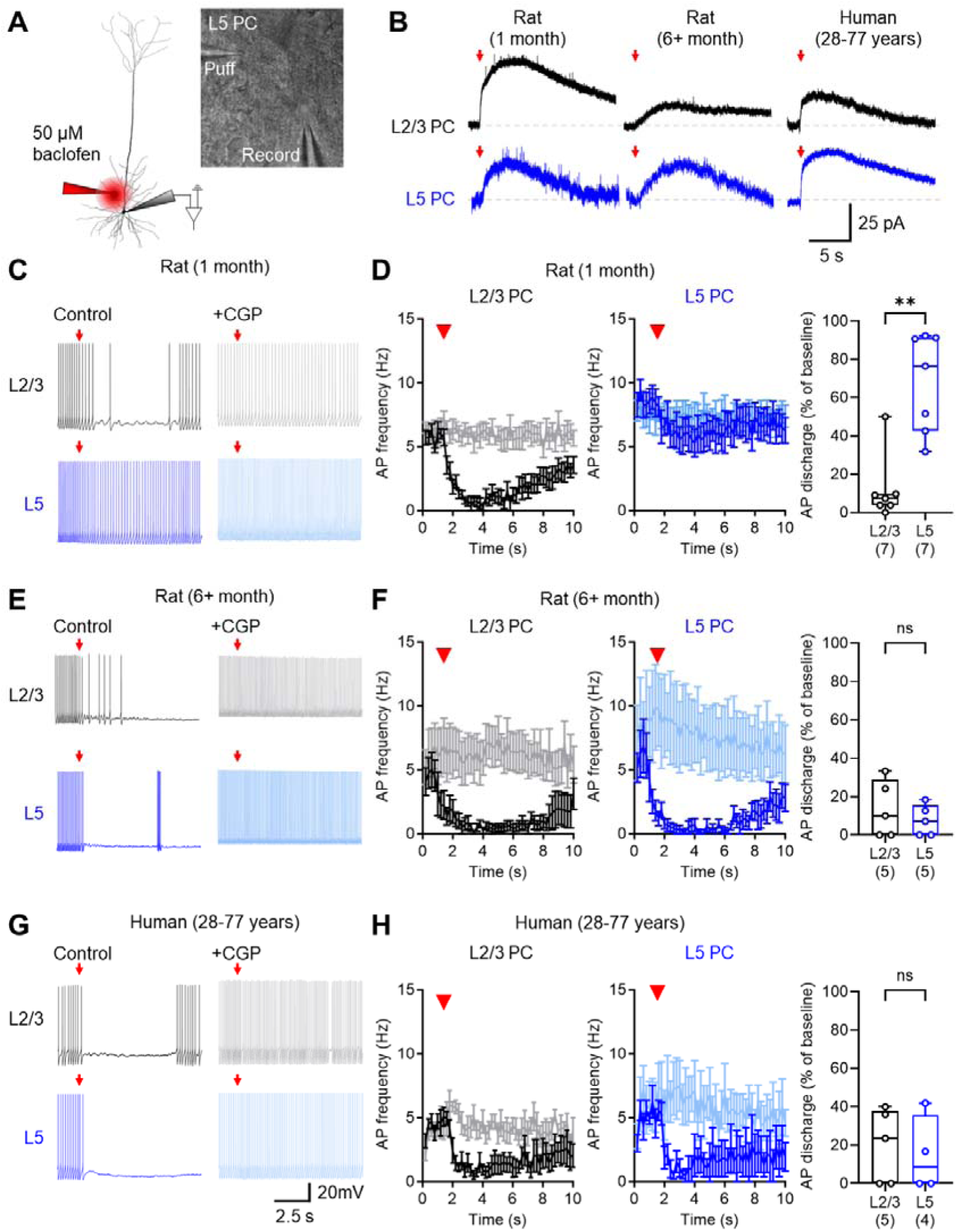
Perisomatic GABA_B_Rs differentially inhibit AP discharge in L2/3 and L5 PCs over development. (**A**) Schematic of experimental set-up depicting puff application of 50 μM baclofen (red) to the perisomatic compartment whilst performing a whole-cell recording. Right, IR-DIC image taken during recording confirming location of somatic recording (record) and baclofen puff pipettes to a L5 PC. (**B**) Example baclofen puff-mediated currents recorded at -65 mV voltage-clamp from the soma in the presence of 10 μM NBQX, 50 μM AP-5 and 10 μM Gabazine, from L2/3 (black) and L5 PCs (blue), from rats at 1 month and 6+ months, or human neurons. Baclofen puff (red arow) and baseline (grey dashed line) are shown for reference. (**C**) Action potential (AP) output of L2/3 (black, upper) and L5 (blue, lower) PCs from 1 month old rats when held at +25 pA above rheobase when subjected to perisomatic baclofen puff under control conditions (left) or in the presence of 5 μM CGP-55,845 (CGP). (**D**) AP output over the duration of puff-recording (200 ms bins) for L2/3 (left) and L5 PCs (middle), compared to AP discharge in the presence of CGP (grey, light blue). Measured change in AP discharge at 2-3 seconds after baclofen puff (right, U _(7,7)_=2, p=0.0023, Mann-Whitney test). Number of recorded cells is shown in parenthesis. (**E, F**) The same analysis, but performed in 6+ month rats. No difference was observed in AP discharge between layers (U _(5,5)_=10, p=0.651, Mann-Whitney test). (**G, H**) No difference was observed in AP discharge between layers in adult human cortex (U _(5,4)_=9, p=0.873, Mann-Whitney test). Data is shown as either mean ± SEM (left and middle panels) or 25 - 75% boxplot with median indicated ± range; all with individual cell data overlain.

### GABA_B_Rs more strongly inhibit presynaptic glutamate release in human cortex, compared to rodents

At glutamatergic synapses, presynaptic GABA_B_Rs rely on heterosynaptic spill-over from neighbouring GABAergic afferents, thus providing powerful circuit-wide control of neurotransmitter release (Kulik *et al*., 2018; Urban-Ciecko *et al*., 2015). We next quantified the magnitude of GABA_B_R presynaptic inhibition at major glutamatergic inputs to L2/3 and L5 pyramidal cells in human and rat cortex. To achieve this, we performed whole-cell patch clamp recordings from L2/3 and L5 PCs at -70 mV voltage-clamp in the presence of picrotoxin (50 μM). We evoked excitatory postsynaptic synaptic currents (EPSCs) in neurons by stimulating L1 (Figure 5A) or L4 (Figure 5C) for afferents to L2/3 PCs, or L1 (Figure 5B) and L5 (Figure 5D) for afferents to L5 PCs, using pairs of stimuli (50 ms interval) to assess the paired-pulse ratio (PPR). To assess the effect of GABA_B_Rs on evoked EPSCs, 10 µM baclofen was applied to the bath for 5 minutes, followed by 5 μM CGP-55,845. These experiments were also performed in 1 month (**Supplementary Figure 2**) and 6+ month old rats (**Supplementary Figure 3**).

**Figure 5:**
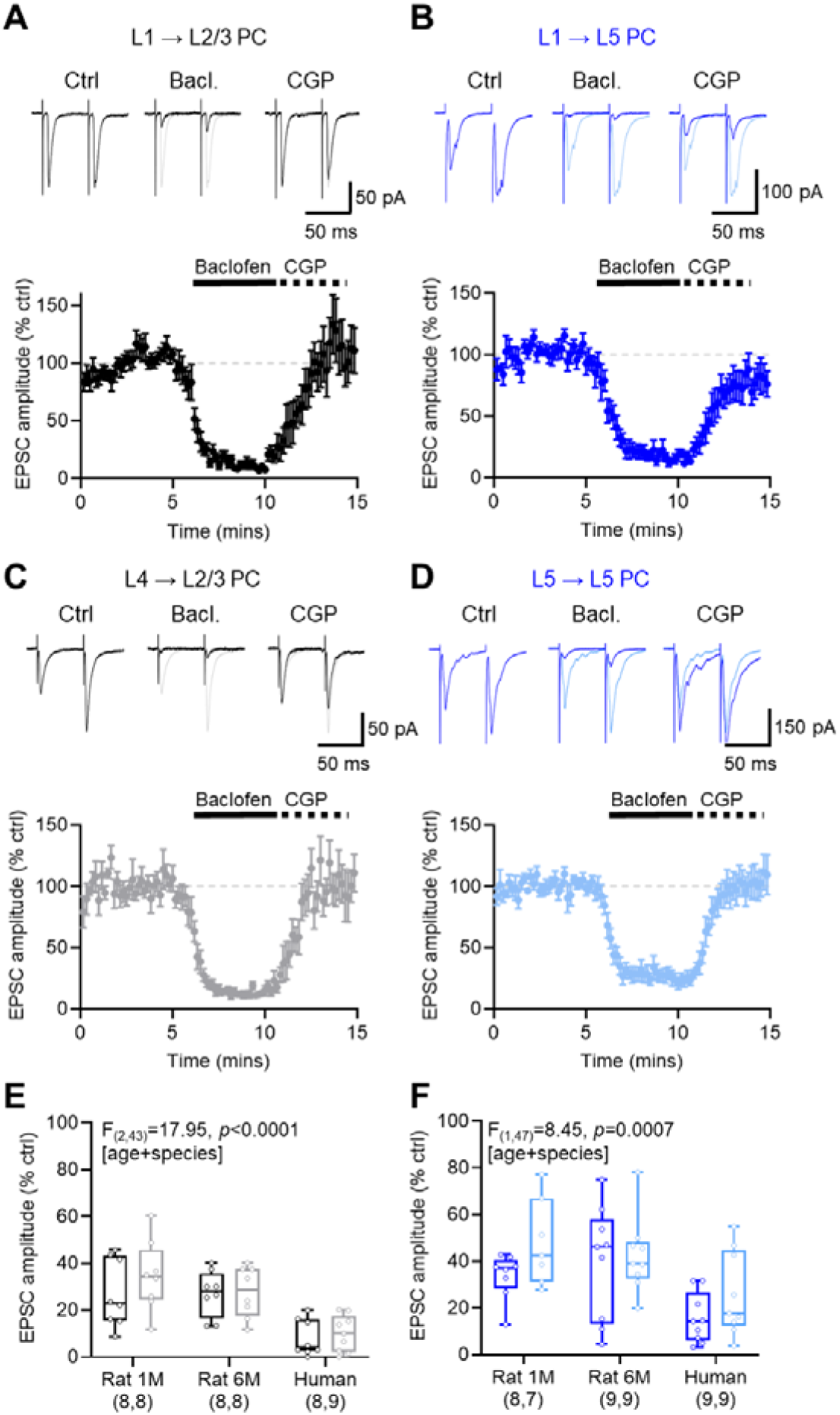
Presynaptic GABA_B_Rs strongly inhibit synaptic inputs to human cortical neurons. (**A**) Example EPSCs evoked by L1 stimulation in L2/3 PCs recorded at -70 mV voltage clamp under control conditions (Ctrl) and following bath application of 10 μM baclofen (Bacl.) and 5 μM CGP-55,845 (CGP). Control recordings are shown for reference (grey traces). Lower, time-course of EPSC amplitude following baclofen (solid bar) and CGP (dashed bar) wash-in. Control baseline is shown for reference (grey dashed line). (**B**) Data in the same form but for L1 inputs to L5 PCs (blue). (**C**) Data in the same form, but for L4 inputs to L2/3 PCs (light grey). (**D**) Data in the same form, but for L5 inputs to L5 PCs (light blue). (**E**) Quantification of baclofen-mediated inhibition of L1 to L2/3 (black) and L4 to L2/3 (grey) inputs to L2/3 PCs during the last minute of wash-in from rats at 1 month (1M), 6+ month (6M) and human neurons. (**F**) The same as (**E**) but L1 to L5 (blue) and L5 to L5 (light blue) inputs to L5 PCs. The number of cells is shown below bars in parenthesis for L1 (black) and L4 (grey) inputs. Data is shown as mean ± SEM (A, B, C, D) or as 25 - 75% boxplot with median indicated ± range. All statistics shown from 2-way ANOVA.

In human L2/3 neurons, stimulation of L1 and L4 produced large amplitude EPSCs which displayed consistent facilitating paired-pulse ratios (PPR; **Supplementary Figure 4**). Bath application of baclofen resulted in a very strong reduction of EPSCs at both L1 and L4 inputs, which was fully blocked by CGP application. Similarly, in L5 PCs we observed a near complete inhibition of L1 and L5 inputs following the baclofen wash-in, which was recovered by CGP application. When we performed the same recordings in 1-month and 6+ month rats we also observed strong reductions in EPSC amplitude following baclofen application, but which was lower than recordings in L2/3 of the human cortex (Figure 5E). Likewise, we found that presynaptic GABA_B_R mediated inhibition in human L5 afferents was stronger than for rats, irrespective of afferent type (Figure 5F).

As presynaptic GABA_B_R activation decreases voltage-dependent Ca^2+^ influx to axon terminals, it can modulate presynaptic release probability. Interestingly, we found that basal release probability of presynaptic afferents displayed path-specific species differences, with L4 stimulation revealing the greatest differences (**Supplementary Figure 5**). With respect to GABA_B_R activation, most synaptic inputs to L2/3 and L5 PCs displayed increased PPR, and thus reduced release probability, upon baclofen application (**Supplementary Figure 4**). Such PPR changes were most prominent at human synapses, especially onto L2/3 PCs – consistent with enhanced presynaptic inhibition. Subsequent CGP application typically blocked baclofen-dependent PPR increases. Quantification revealed that PPR was more strongly modulated by baclofen at inputs to human L2/3 PCs than for either age of rat, an effect that was not observed for L5 inputs (**Supplementary Figure 5**). Taken together, these data show that GABA_B_R presynaptic inhibition is stronger in presynaptic afferents terminating onto human cortical neurons. Given the greater overall inhibitory tone, this suggests that presynaptic GABA_B_Rs may strongly inhibit active human cortical circuits.

### Population oscillations in human cortex are highly sensitive to GABA_B_R activation

So far, we have shown that GABA_B_Rs can strongly inhibit both pre- and postsynaptic domains of L2/3 and L5 PCs of the adult human cortex. As endogenous GABA_B_R activation requires heterosynaptic spill-over of GABA (Scanziani, 2000), the effects of GABA_B_R activation may be most prominent in active circuits (Urban-Ciecko *et al*., 2015). To determine the circuit-level effects of GABA_B_R activation, we conducted local field potential (LFP) recordings to assess how *in vitro* neuronal oscillations are affected by baclofen. Persistent neuronal oscillations were pharmacologically induced using kainic acid (KA, 0.6 - 2 µM) and carbachol (CCh, 60-200 µM), as previously described in rodent (Buhl et al., 1998; Cunningham et al., 2003; Johnson et al., 2017) and human cortical tissue (Pennifold, 2017). Following 30-40 minute KA and CCh application we reliably observed broadband neuronal oscillations in L2/3 and L5 of human neocortex (Figure 6A), with Fast-Fourier transform displaying prominent oscillations in theta-alpha (4-10 Hz), beta (10-30 Hz), low-gamma (30-50 Hz), and high-gamma (51-85 Hz) frequency bands (Figure 6B). Similar oscillatory activity was also observed in simultaneous recordings from L5 (Figure 6C). We found no overt difference in the strength of KA+CCh induced oscillations between L2/3 and L5 (Figure 6D).

**Figure 6:**
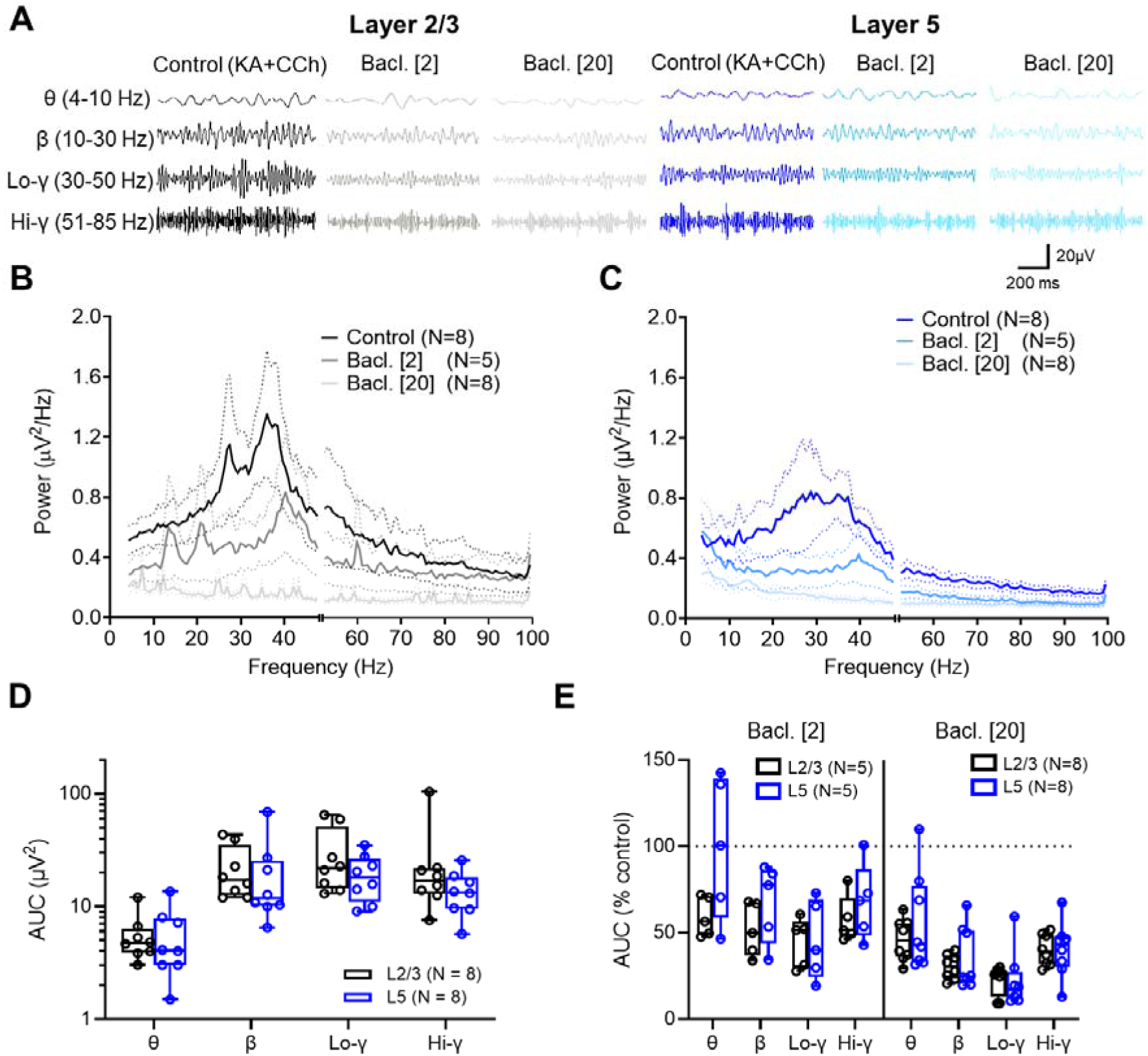
GABA_B_R activation strongly attenuates oscillatory activity in L2/3 and L5 of the human cortex. (**A**) Example Local field potential (LFP) signals band-pass filtered into theta (θ), beta (β), low-gamma (Lo-γ), or high-gamma (Hi-γ) ranges of frequencies; in L2/3 (black) and L5 (blue). Data are shown during control (Kainic Acid, KA [0.6 - 2 µM] + Carbachol, CCh [60-200 µM]) and during the last 2 minutes of 2 µM (light blue and grey) or 20 µM (lightest blue and grey) baclofen bath application. (**B**) Power spectra from Fast-Fourier transforms of filtered L2/3 LFP recordings under control oscillatory conditions (black) and in the presence of 2 µM (grey) or 20 µM (light grey) baclofen (Bacl.). (**C**) Data shown in the same format but from L5 under control conditions (blue) and following 2 µM (light blue) and 20 µM (lightest blue) baclofen. (**D**) Integrals of each frequency epoch, shown as area-under-curve (AUC), for L2/3 (black) and L5 (blue) LFP recordings revealed no difference in overall power (F_(1, 14)_ = 0.99, *p*=0.336, 2-way ANOVA [layer]). (**E**) Comparison of the effect of baclofen on oscillatory power for L2/3 (black) and L5 (blue) LFP recordings. Data is shown as the % change from control AUC for each frequency band, with respect to either 2 μM (left) and 20 μM (right) baclofen, with statistical effects of baclofen concentration (F_(1, 22)_ = 19.2, *p*=0.0002 [baclofen concentration], 3-way ANOVA), and layer-wise inhibition (F_(3, 66)_ = 3.782, *p*=0.014 [layer], 3-way ANOVA). Data are shown as mean ± SEM (B, C), or 25 - 75% boxplot with median indicated ± range; with individual data point overlain.

GABA_B_R activation has been shown to abolish theta and gamma oscillations in rodent brain slices (Booker *et al*., 2020; Brown *et al*., 2007) and with complex actions on human EEG traces (Badr et al., 1983). Following generation of stable oscillations, we first applied baclofen at 2 μM, a low concentration likely to impact presynaptic receptors only (Dugladze et al., 2013). In both L2/3 and L5, this led to a large reduction in oscillatory power over the range of frequencies, tested 30-40 minutes after application (Figure 6E). Subsequent bath application of 20 μM baclofen, a saturating concentration, produced further large reductions in oscillatory power at all frequencies. Comparing the effect of baclofen between recordings, we found that the degree of baclofen inhibition on oscillatory power was less prominent in L5 than L2/3.

We performed equivalent experiments in L2/3 of 6+ month rats (**Supplementary Figure 6**), whilst the overall strength of oscillations in was comparable between species when using the same KA and CCh pharmacology, the degree of inhibition mediated by bath application of baclofen was notably lower in rat L2/3. Despite minimal difference of postsynaptic GABA_B_R function in L2/3 of humans and rats, this may suggest that presynaptic GABA_B_Rs play a dominant role in cortical oscillations. Together, these data show that GABA_B_R activation strongly controls the strength of oscillatory activity in human cortex.

### GABA_B_R activation reduces the synchrony of cortical oscillations

Distinct neuronal oscillations interact with and synchronise each other, a process termed cross-frequency coupling (Canolty and Knight, 2010), where neuronal firing patterns align to produce coordinated circuit-level activity. We found low concentrations of baclofen to inhibit the strength of oscillations across a broad range of frequencies, likely due to the activation of presynaptic GABA_B_Rs, we next asked if their activation leads to reduced correlation of different oscillatory frequencies. For this, we performed more in-depth analysis of LFP recordings and how oscillations interact (Figure 7A). First, autocorrelation analysis revealed that each of the oscillatory bands were highly consistent for both L2/3 and L5 (Figure 7B). Next, we performed cross-correlation analysis between different frequency bands, notably theta vs. beta, theta vs. low-gamma, beta vs. low-gamma, and beta vs. low-gamma, under control conditions and following 2 μM and 20 μM baclofen in L2/3 (Figure 7C). Similar cross-correlations were observed in L5 (**Supplementary Figure 7**). In L2/3, we found that baseline correlation strength varied widely between frequency pairs (F_(3, 28)_ = 14.95, P<0.0001, 2-way ANOVA [frequency]), and which was highly sensitive to baclofen application (F_(1.278, 28.12)_ = 29.18, 2-way ANOVA [baclofen], Figure 7D). For L5, we also observed high variability between frequency pairs (F_(3, 28)_ = 17.05, P<0.0001, 2-way ANOVA [frequency]), and which was highly sensitive to baclofen application (F_(2, 44)_ = 90.07, 2-way ANOVA [baclofen], Figure 7E). We found that there was minimal synchrony between L2/3 and L5 when cross-correlation was performed in a frequency specific manner between layers, while some slices showed prominent correlation, in particular in the low-gamma range, they were not affected by baclofen application (**Supplementary Figure 8**).

**Figure 7:**
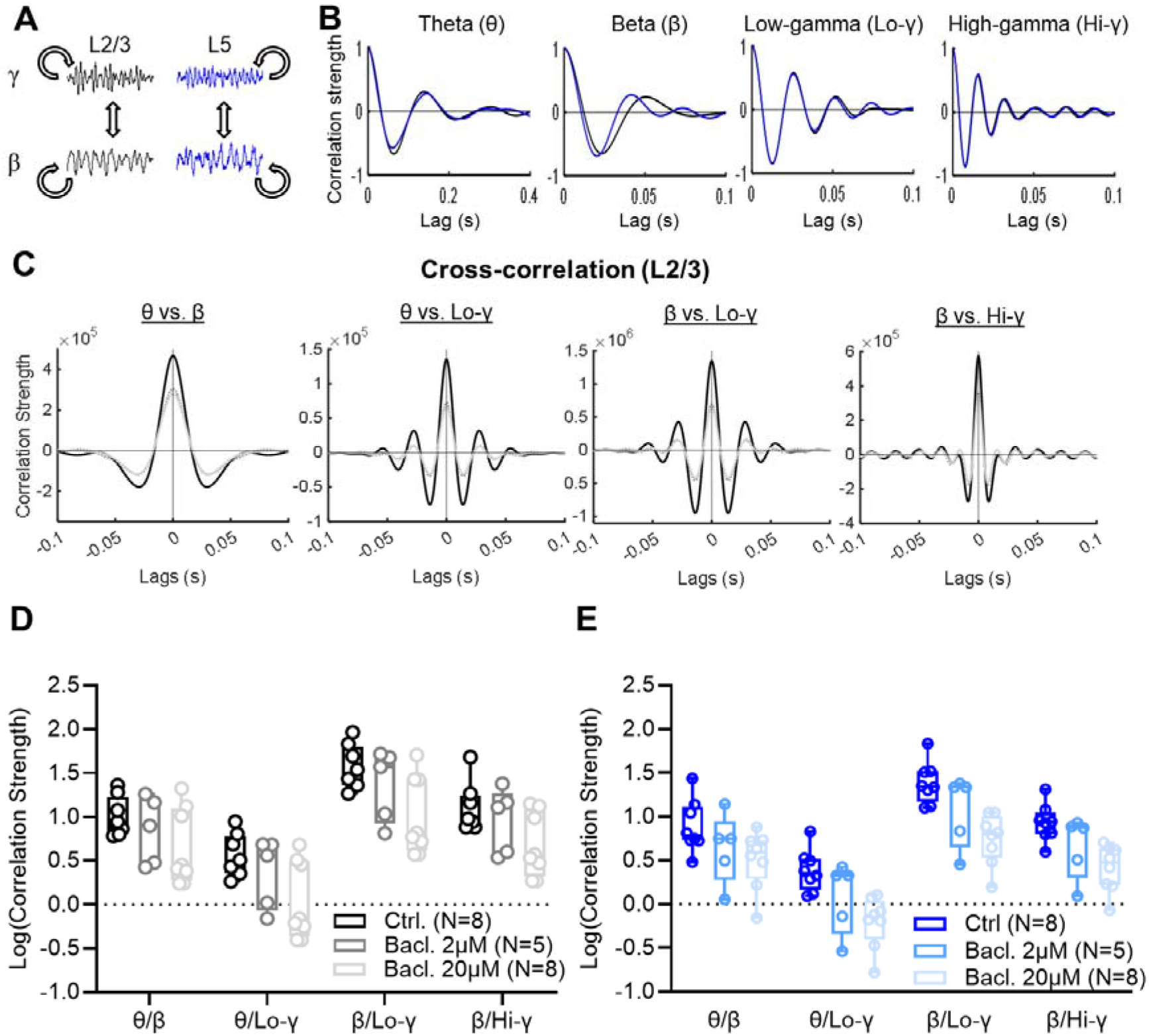
GABA_B_R activation de-synchronises beta/gamma coupling in human cortical circuits. (**A**) Schematic of auto-correlation and cross-correlation analysis strategy. (**B**) Example plots of auto-correlation strength as a function of lag-time (ms) in LFP recordings from L2/3 (black) and L5 (blue) of the human cortex under control conditions (Kainic Acid, KA [0.6 - 2 µM] + Carbachol, CCh [60-200 µM]). (**C**) Example plots of cross-correlation strength between prominent oscillations in LFP recording from human L2/3 under control conditions (black) and following 2 µM (dark grey) or 20 µM (light grey) baclofen bath application. (**D**) Comparison of log-transformed correlation strength at zero lag (no time delay) between key oscillatory bands in L2/3 revealed a significant frequency (F_(3, 28)_ = 14.95, P<0.0001, 2-way ANOVA [frequency]), and baclofen effect (F_(1.278, 28.12)_ = 29.18, 2-way ANOVA [baclofen]. (**E**) Similar observations on cross-correlation strength were observed in human cortex L5 with respect to frequency (F_(3, 28)_ = 17.05, P<0.0001, 2-way ANOVA [frequency]) and baclofen application (F_(2, 44)_ = 90.07, 2-way ANOVA [baclofen]. All data is shown as 25 - 75% boxplot with median indicated ± range; with individual values overlain.

Finally, we examined phase-amplitude coupling (PAC) between these oscillation frequency pairs. PAC represents one form of cross-frequency correlation, wherein the phase of a lower frequency oscillation modulates the amplitude of a higher frequency (Canolty and Knight, 2010). PAC plays a crucial role in orchestrating coordinated local and global neuronal activity across spatial and temporal scales (Jensen and Colgin, 2007; Tort et al., 2010); especially theta/low-gamma and beta/low-gamma, which are associated with memory encoding (Lisman and Jensen, 2013) and sensorimotor function (Yanagisawa et al., 2012). We found that theta vs. low-gamma and beta vs. high-gamma oscillatory pairs possessed prominent phase-amplitude coupling, but this was not affected by the application of baclofen (**Supplementary Figure 9**).

These data indicate that GABA_B_Rs contribute to the synchronous activity of local cortical circuits, including the interactions between oscillatory frequencies. Given the role of somatostatin and parvalbumin interneurons (Cardin et al., 2009; Chen et al., 2017) in the generation of beta and gamma oscillations, it is plausible that such loss of inter-frequency correlation indicates the possibility of cell-type specific effects. As perturbations in PAC strength have been linked to cognitive impairments, such as in learning and memory (Lisman and Jensen, 2013) the loss of beta and low-gamma synchrony indicates that even modest concentrations of baclofen may impair cortical circuit function *in vivo*.

### Clinical presentation of seizures is correlated with higher GABA_B_R activation

A key consideration of human resected brain tissue is the seizure history (Dührsen et al., 2019), which may have an impact on GABA_B_R expression (Princivalle *et al*., 2003; Rocha et al., 2015; Sheilabi *et al*., 2018; Teichgräber *et al*., 2009). So far, we have only presented data from patients who did not experience seizures in their recent clinical history, however tissue was also collected from 15 patients who had experienced seizures and been prescribed Levetiracetam (LEV, 1g/day) at least 3 days prior to surgery. In addition, 3 patients had been prescribed LEV, but had not experienced seizures. We interrogated our data to ask what effect a recent history of seizures and/or LEV treatment may have on neuronal activity and GABA_B_R signalling. For this, we compared this data to that collected under identical circumstances to L2/3 neurons from seizure-free individuals (**Supplementary Table 1**).

Consistent with administration of acute LEV (Englund et al., 2011), we observed no difference in AP discharge responses in L2/3 PCs from individuals who had received LEV (Figure 8A), nor in the ability of neurons to repetitively fire trains of APs (Figure 8B). Assessing GABA_B_R function, we first examined slow-IPSCs resulting from L1/2 stimulation (as in Figure 3). We found no overt change in GABA_B_R-mediated IPSC amplitudes in patients with Seizures/LEV (20% greater, t_(19,19)_=0.41, p=0.68, Holm-Sidak test), but individuals who had received LEV, but not experienced seizures had slow-IPSCs 342% larger than controls (t_(19,5)_=3.2, p=0.005, Holm-Sidak test; Figure 8C). Consistent with larger slow-IPSCs, we found that wash-in of baclofen revealed greater whole-cell currents in patients receiving LEV (irrespective of seizure history) than that of LEV-free individuals (Figure 8D). Quantification of current-density revealed that patients who had experienced seizures had 59% greater baclofen-mediated currents than Seizure/LEV-free patients (t_(24,19)_=3.3, p=0.003, Holm-Sidak test), with LEV/Seizure-free patients having greater GABA_B_R current densities, 95% higher than control patients (t_(19,5)_=3.3, p=0.0034, Holm-Sidak test; Figure 8E). To determine whether acute LEV increases GABA_B_R currents, we performed a subset of experiments in which human brain slices were incubated with 100 μM LEV for 3 hours prior to recording. We found that in brain slices from control patients and those who had received LEV prior to surgery, subsequent *in vitro* application of LEV did not lead to enhanced baclofen-mediated currents, despite overall patient group differences (Figure 8F).

**Figure 8:**
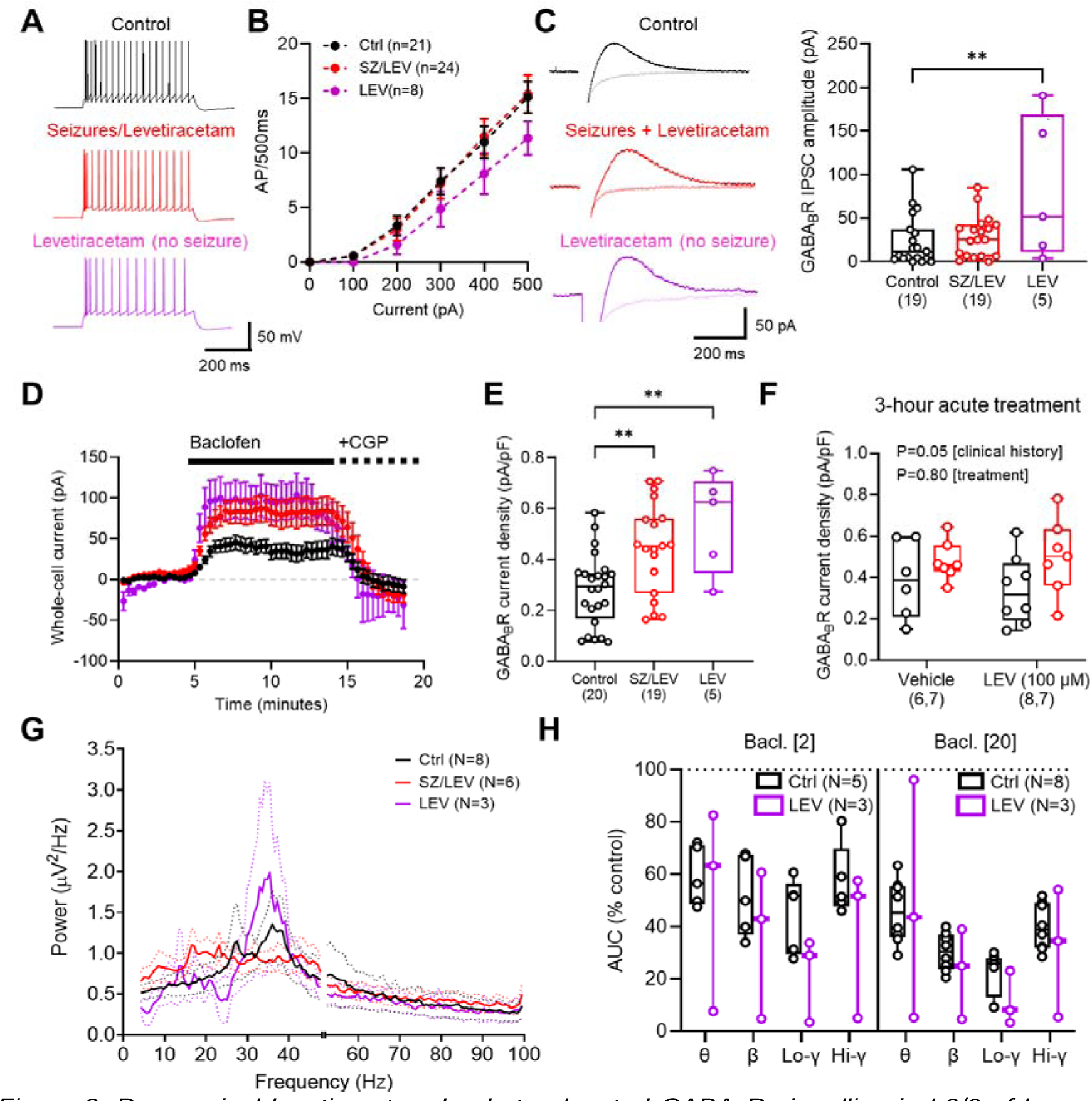
Pre-surgical levetiracetam leads to elevated GABA_B_R signalling in L2/3 of human neocortex. (**A**) Example responses of human L2/3 PCs to 500 pA depolarising current injections, from seizure-free control patients (black, upper), patients who have experienced seizures and received levetiracetam (red, SZ&LEV, middle), or patients who did not experience seizures, but received levetiracetam (purple, LEV, lower). (**B**) Current*-*frequency response plots (0 – 500 pA, 500 ms), from control (21 cells from 12 cases), SZ&LEV (24 cells from 11 cases), and LEV (8 cells from 3 cases), revealed no overt effect on AP discharge (F_(2, 50)_ = 0.77, *p*=0.47, 2-way repeated measures ANOVA). (**C**) Example slow-IPSCs evoked in the presence of 10 μM NBQX, 50 μM AP-5, 10 μM gabazine following 5x 200 Hz (50 V) stimulation to the L1/2 border in control (black), SZ&LEV (red), and LEV (purple), which are superimposed over the same response recorded in the presence of 5 μM CGP-55,845 (CGP, light traces). Quantification of slow-IPSC amplitudes for all recorded cells from each patient group (F_(2, 40)_= 5.3, *p*=0.009, 1-way ANOVA). (**D**) Time-course of whole-cell current changes following 10 μM baclofen (black bar) and then CGP (dashed bar) for each patient group. (**E**) Quantification of baclofen-mediated current density for cells from each group (F_(2, 45)_ = 8.6, *p*=0.007, 1-way ANOVA). (**F**) Baclofen-mediated current density for slices pre-treated for 3 hours with 100 μM levetiracetam (LEV) or vehicle, from control and SZ&LEV patients. No effect of acute levetiracetam was noted on baclofen current density (F_(1, 25)_ = 0.06, *p*=0.81, 2-way ANOVA [treatment]), but LEV&SZ current density remained higher than control (F_(1, 25)_ = 4.55, *p*=0.043 [patient group]). (**G**) Power spectra from LFP recordings of L2/3 in the presence of KA+CCh from control (8 cases), SZ&LEV (6 cases), and LEV (3 cases) patients. (**H**) Quantification of change in AUC from LEV patients (3 cases) compared to control recordings, in the presence of 2 μM (left) or 20 μM (right) baclofen. Prior history of LEV treatment was significantly different from control (F_(1, 60)_ = 6.64, *p*=0.012, 3-way ANOVA). All data is shown as either mean ± SEM (B, D, G) or 25 - 75% boxplot with median indicated ± range (C, E, F, H) ; with individual values overlain. All statistics from 1, way (C, E), 2-way (F) or 3-way (H) ANOVA, with Holm-Sidak tests (as appropriate).

To ascertain whether experiencing seizures and administration of LEV altered circuit function, we assessed oscillatory activity following KA+CCh application. We found no overt difference in oscillatory power between patients, irrespective of clinical background (Figure 8G). As patients who had not experienced seizures, but had been prescribed LEV, displayed the greatest GABA_B_R-mediated signalling (compared to control), we asked how baclofen effected oscillations in brain slices from these patients. Comparison of the change in power, as measured as % difference from control of the AUC, revealed that patients who had experienced seizures and received LEV displayed a greater reduction in circuit activity in response to baclofen than control patients independent of oscillatory frequency (Figure 8H).

These data show that LEV boosts endogenous GABA_B_R signalling, through a mechanism that occurs over long (>3 hour) timescales. This increase in GABA_B_R signalling enhances the baclofen sensitivity of neuronal oscillations, which may underlie the anti-seizure properties of LEV itself.

## Discussion

In this study we provide the first quantitative assessment of functional GABA_B_R control of human brain circuits, directly comparing this to that of adult and juvenile rats. We show that in adult human brain slices, postsynaptic GABA_B_Rs produce robust currents in L2/3 and L5 PCs which are largely stable over adult life. In contrast, GABA_B_R-mediated currents in rodent neurons displayed a rapid age-dependent decline from adolescence into adulthood. Conversely, presynaptic GABA_B_R mediated inhibition is largely stable across rodent development, but is weaker that human cortical circuits. These functional GABA_B_Rs control both the strength and timing of neuronal oscillations in living human brain circuits, which are potentially more sensitive to modulation of presynaptic GABA_B_Rs when compared to adult rodent circuits. Finally, we show that the anti-seizure medication levetiracetam boosts endogenous GABA_B_R levels, in a manner that is independent of seizure status. These data provide a mechanism by which GABA_B_Rs contribute to the long-term prevention of seizure activity and epilepsy in human brain circuits.

### Differential GABA_B_R mediated control of neuronal excitability across life-span and species

Our data shows that human neurons possess strong and reliable postsynaptic GABA_B_R signalling, regardless of location within the cortical column. Our data extends earlier studies that confirmed the presence of GABA_B_Rs in adult human cortical tissue (Deisz, 1999; Teichgräber *et al*., 2009). We significantly expand these findings, by performing comparative analysis with rodents, across layers, and with respect to clinical state. This comparison shows that L5 pyramidal cells in juvenile rats largely lack measurable GABA_B_R currents at the soma, which agrees with earlier studies in young rats (<2 months old) that GABA_B_Rs are confined to the distal dendritic tuft (Schulz *et al*., 2021) and contribute to inter-hemispheric inhibition of neuronal activity (Palmer *et al*., 2013; Palmer et al., 2012). While such specific dendritic processes are likely to be maintained into adulthood, the output of cortical columns from L5 is likely to be under stricter control of slow inhibitory signalling as we age. Evidence for such age dependent changes in GABA_B_R expression are scant, however it is known that baclofen has an age-dependent effect on tonic-clonic seizures in rats (Velíšková et al., 1996), which may provide evidence of such changes.

We also observed that GABA_B_R mediated presynaptic inhibition of glutamate release was strongest in human tissue, compared to rats at any age tested. In line with a role of presynaptic GABA_B_Rs in controlling circuit activity (Urban-Ciecko *et al*., 2015) and seizure activity (Dugladze *et al*., 2013; Mangan and Lothman, 1996), we observed that low (likely presynaptic selective) concentrations of baclofen reduced the power of oscillations human and rat brain slices. Together, these data strongly suggest that glutamatergic synapses onto human neurons are under greater presynaptic inhibitory control. Given that lower concentrations of baclofen are required for presynaptic activation, this indicates that clinical administration of low dose baclofen may have central effects, including sedation (Meythaler et al., 2004) and impaired memory (McDonnell et al., 2007). In rodents, electrical stimulation suppressed KA-induced gamma oscillations on a timescale matching GABA_B_R-mediated slow IPSPs (Brown *et al*., 2007), suggesting that the pro-cognitive effects of GABA_B_R antagonism (Getova and Bowery, 1998; Lasarge et al., 2009) may be due to suppression of gamma oscillations. Such a strong pre- and postsynaptic function of GABA_B_Rs to the perisomatic domains of L2/3 and L5 PCs in humans could explain why GABA_B_R modulation has enhanced effects on learning and memory as we progress through life (Lasarge *et al*., 2009).

We show that while L1/2 stimulation produced GABA_B_R-mediated IPSCs in rats and humans, perisomatic GABA_B_Rs exert large functional effects on neuronal AP discharge. A number of studies have confirmed that GABAergic terminals in upper cortical layers can activate distal dendritic GABA_B_Rs (Hay et al., 2021; Oláh et al., 2007; Palmer *et al*., 2012; Pérez-Garci *et al*., 2013; Schulz *et al*., 2021; Urban-Ciecko *et al*., 2015). However, these likely minimally contribute to perisomatic inhibition in deep L2/3 and L5. The source of GABA for these receptors remains elusive, but likely originates from spillover of GABA from various synapses (Booker *et al*., 2013; Oláh *et al*., 2007; Scanziani, 2000; Urban-Ciecko *et al*., 2015) leading to entrainment of PCs to local activity. Indeed, we show compelling evidence that GABA_B_Rs modulate cortical oscillations, where circuit wide activation of GABA_B_Rs reduce neuronal discharge and desynchronises oscillatory activity. Beyond fast neuronal oscillations, high densities of GABA_B_Rs on L5 neurons could contribute to the timing of slow-wave oscillations and the up-down states of these neurons (Craig et al., 2013).

### The role of GABA_B_Rs contributing to human brain health and pathology

GABAergic neurotransmission has long been linked to the onset and development of seizure disorders, albeit the role of ionotropic GABA_A_Rs display complex and paradoxical effects on seizure propagation (Ben-Ari, 2006; De Curtis and Avoli, 2016), with many anti-seizure medications targeting these receptors and GABA release (Richardson et al., 2024). GABA_B_Rs have received much less attention as potential therapeutic targets for seizures, due to the fact that the agonist baclofen can exacerbate seizures (Kofler et al., 1994), and antagonists have bidirectional effects (Mareš and Šlamberová, 2006). This complexity is likely due to potent GABA_B_R signalling in both excitatory and inhibitory neurons alike, leading to shifts in excitatory/inhibitory balance that may produce complex circuit effects (reviewed in (Kulik *et al*., 2018)).

GABA_B_Rs themselves have long been implicated in seizure disorders, including temporal lobe epilepsy (TLE), where expression of receptor subunits has been suggested to be elevated in post-mortem human brain (Princivalle *et al*., 2003; Sheilabi *et al*., 2018), but with impaired function in neurons recorded from the seizure focus (Teichgräber *et al*., 2009). This is mimicked in rodent models of TLE, where rapid loss of GABA_B_R subunits was observed in dorsal hippocampus (Straessle *et al*., 2003). Our data does not address whether GABA_B_Rs are lost following seizures, however, we show that in patients who have received LEV functional GABA_B_R currents are elevated, irrespective of seizure history. The mechanism by which LEV mediates long-term suppression of seizures is not fully understood. It is known that it directly binds to Synaptic Vesicle Protein 2A (SV2A) (Lynch et al., 2004), which is present in axon terminals of both glutamatergic and GABAergic neurons, and may be strongly expressed in basket cell axon terminals (Vanoye-Carlo and Gómez-Lira, 2019). Indeed, administration of LEV leads to increased GABA levels in the brain (Doelken et al., 2010), which agrees with a central role at GABAergic synapses. The mechanisms by which LEV leads to elevated GABA_B_R currents remains to be determined. However, this is likely a specific effect, as previous studies of GABA_A_R signalling in resected human brain tissue did not observe an effect on ionotropic currents (Field et al., 2021). The ability to regulate endogenous GABA_B_R functional currents represents a unique and promising avenue for the development of future anti-seizure medications, which may lack many of the deleterious side effects of LEV (French et al., 2001; Halma et al., 2014)

### Technical considerations

Given the nature of the recordings we have performed, there are several key technical considerations. First, all intracellular recordings performed in the current study were made from the cell body. This could largely overlook the role of GABA_B_Rs in the distal dendrites, due to electrotonic compartmentalisation, particularly for L5 neurons (Beaulieu-Laroche et al., 2018; Palmer *et al*., 2012; Schulz *et al*., 2021). While we fully appreciate the importance of dendritic GABA_B_Rs, particularly in the case of integration of synaptic inputs, the purpose of this study was to assess the impact on the spiking ability of neurons in response to activity within their layer. Determining how GABA_B_Rs contribute to dendritic computation in human neurons is a worthwhile undertaking, especially given differences in dendritic arborisation and electrotonic propagation (Kalmbach *et al*., 2018), but is out with the scope of this current study.

Second, our human data has been collected from a relative old patient cohort (Median age: 60 years). We sought to address this age of patients by performing parallel experiments in adult rats, but there still remains a clear chronological difference in brain age between these groups (Sengupta, 2013), which may make direct comparison complicated, which is limited by the patients we can obtain brain samples from. Furthermore, our human cohort does not fully capture adult brain development, as our youngest sample was 28 years old. Further studies examining more adolescent or paediatric cohorts would address this issue. Nevertheless, understanding the function of GABA_B_Rs across the adult lifespan into old-age has distinct merits, in particular as clinical trials for baclofen are often in older patient cohorts (Garbutt et al., 2010), GABA_B_R expression may display age dependent effects (Pandya *et al*., 2019) and GABA_B_Rs are implicated in neurodegenerative conditions (e.g. Alzheimer’s disease (Chu et al., 1987)).

## Conclusion

In conclusion, we provide the first quantitative analysis of functional GABA_B_R mediated currents, and their role in controlling cellular and circuit level excitability in the adult human cortex. We show that humans display phylogenetic differences in the functional control of neuronal activity at both presynaptic and postsynaptic GABA_B_Rs, which efficiently control local circuit activity, and are crucially involved in the mechanism of action of the anti-seizure medication levetiracetam. These data highlight the importance of GABA_B_R signalling mechanisms over the lifespan of humans, particularly with respect to epilepsy research and the generation of new anti-seizure medications.

## Acknowledgments

Most of all we would like to thank the patients who generously agreed to donate tissue for this study. NHS Lothian NRS BioResource and Tissue Governance unit and EMERGE Research Nurse team, in particular Allan MacRaild, Ikeoluwa Adekoya, Anuka Boldbaatar and Sarah Risbridger. Thanks are given to David JA Wyllie and Peter C Kind for the support to conduct this work. We also thank Naima Elose Borras for help with initial image analysis; Kind Lab members for helpful discussions; and Colin Smith, Mark Cunningham, and Giles Hardingham for initial discussions around establishing tissue collection. Finally, this project was funded through generous support from the Simons Initiative for the Developing Brain (SAB, MW), Edinburgh Neuroscience Neuroresearchers Fund (SAB), RS McDonald Seedcorn Fund (SAB), Medical Research Council UK (FLB), Race Against Dementia (CD), The James Dyson Foundation (CD), Alzheimer’s Society UK (CD)

## Author contributions

**Max Wilson:** Investigation, Methodology, Software, Formal analysis; Visualization, Writing - Original Draft, Writing - Review & Editing**; Lewis W. Taylor** - Resources, Writing - Review & Editing; **Soraya Meftah** - Resources, Writing - Review & Editing, **Robert McGeachan** - Resources, Writing - Review & Editing; **Tamara Modebadze**: - Resources, Writing - Review & Editing; **Ashan Jayasekera:** Resources; **Christopher Cowie:** Resources; **Fiona Lebeau**: Resources, Project administration; **Imran Liaquat**: Resources; **Claire Durrant**: Resources, Project administration, Writing - Review & Editing; **Paul Brennan**: Resources Methodology, Project administration, Writing - Review & Editing; **Sam Booker**: Conceptualization, Methodology, Validation, Formal analysis, Visualization, Writing - Original Draft, Writing - Review & Editing, Project administration, Funding acquisition.

## Data availability

All data presented in the figures and text are available in Supplementary Materials. Any further data will be made available upon reasonable request.

## Methods

### Animals

*In vitro* electrophysiological experiments were performed in acute slices from either 28-38 day-old or 6-8 month old male Long-Evans Hooded rats. All experiments were performed in accordance with institutional (University of Edinburgh, UK) and UK Home Office guidelines (ASPA: PPL: P2262369). All rats were maintained in 12 hour light/dark cycles, housed in litter-mate cages of 2-5 rats, and given *ad libitum* access to food and water.

### Resected human tissue collection

All human brain tissue collection was subject to local and regional ethical approval (NHS Lothian: REC number: 15/ES/0094, IRAS number: 165488; NHS Newcastle: IRAS 173990). Prior to tissue collection, all patients provided written consent for access tissue and anonymised patient information usage (Edinburgh: NHS Lothian Caldicott Guardian Approval Number: CRD19080). No patient identifying information was stored and all patient information anonymised. Human neocortical brain tissue was collected during the resection of brain tumours (**Supplementary Table 1**). Based on our data, we identified 3 key human groups: control individuals (no seizure history or anti-seizure medication), recent seizure history & pre-operative levetiracetam, and patients who had received levetiracetam (but who had not experienced seizures). This latter group had been prescribed prophylactic levetiracetam 500 mg twice daily, for at least 3 days prior to surgery (Jenkinson et al., 2019), as part of a now-closed clinical trial (EudraCT Number: 2018-001312-30).

Brain tissue was collected and sliced as previously described (McGeachan et al., 2024; Taylor et al., 2024). Briefly, following surgical opening of the skull and dura, a small piece of cortical tissue was resected *en route* to accessing more deeply sited tumours. This brain tissue was then rapidly transferred to ice-cold, oxygenated HEPES modified artificial cerebrospinal fluid (HEPES-ACSF; in mM: 87 NaCl, 2.5 KCl, 10 HEPES, 1.25 NaH_2_PO_4_, 25 Glucose, 90 Sucrose, 1 Na-Pyruvate, 1 Na-Ascorbate, 7 MgCl_2_, and 0.5 CaCl_2_; pH 7.35 with NaOH) and then transported to the laboratory (c.a. 20-40 minutes). Brain tissue was then mounted in 3% agar gel (nominally 30 °C), blocked and mounted in an oscillating blade vibratome (VT1200S, Leica, Germany). For whole-cell recordings, 300 µm thick slices of cortex were cut and transferred to either a submerged storage chamber containing sucrose-modified ACSF (in mM: 87 NaCl, 2.5 KCl, 25 NaHCO_3_, 1.25 NaH_2_PO_4_, 25 glucose, 75 sucrose, 7 MgCl_2_, 0.5 CaCl_2_, 1 Na-Pyruvate, 1 Na-Ascorbate) warmed to 35 °C and bubbled with carbogen (95% O_2_/5% CO_2_) for 30 min then placed at room temperature. For extracellular recordings, 500 μm thick slices were cut and stored in a liquid/gas interface chamber containing recording ACSF (in mM: 125 NaCl, 2.5 KCl, 25 NaHCO_3_, 1.25 NaH_2_PO_4_, 25 glucose, 1 MgCl_2_, 2 CaCl_2_); bubbled with carbogen and maintained at room temperature.

### Rat brain slice preparation

Rat brain slices containing primary somatosensory cortex were prepared as previously described (Booker, 2020). Briefly, rats were sedated with isoflurane, anaesthetised with sodium pentobarbital, then transcardial perfusion performed with carbogenated (95%O_2_/5% CO_2_) ice-cold sucrose-ACSF. Following perfusion rats were decapitated, and their brain rapidly removed into ice-cold, carbogenated sucrose-ACSF. For whole-cell recordings, 400 μm thick coronal brain slices were cut on a Vibratome (VT1200s, Leica, Germany) in semi-frozen sucrose-ACSF, then stored submerged in sucrose-ACSF warmed to 35°C for 30 min and subsequently at room temperature. For extracellular oscillation analysis, 500 μm thick slices were cut and stored in a liquid/gas interface chamber containing recording ACSF; bubbled with carbogen and maintained at room temperature.

### Intracellular recordings

For whole-cell patch-clamp recordings, slices were transferred to a submerged recording chamber which was perfused with carbogenated recording ACSF (in mM: 125 NaCl, 2.5 KCl, 25 NaHCO_3_, 1.25 NaH_2_PO_4_, 25 glucose, 1 MgCl_2_, 2 CaCl_2_) at 5-6 mL/min at 31 ± 1°C by an inline heater. Slices were visualized under Köhler illumination by means of an upright microscope (Slicescope, Scientifica, UK), equipped with a 40x water-immersion objective lens (N.A. 0.8; Olympus). Whole-cell patch-clamp recordings were accomplished using a Multiclamp 700B amplifier (Molecular Devices, CA, USA). Recording pipettes were pulled from borosilicate glass capillaries (1.5 mm outer/0.86 mm inner diameter, Harvard Apparatus, UK) on a horizontal electrode puller (P-97 or P-1000, Sutter Instruments, CA, USA). When filled with intracellular solution (in mM: 142 K-gluconate, 4 KCl, 0.5 EGTA, 10 HEPES, 2 MgCl_2_, 2 Na_2_-ATP, 0.3 Na_2_-GTP, 10 Na-phosphocreatine, 0.1% Biocytin, corrected to pH 7.4 with KOH, 295–305□mOsm) a pipette resistance of 2-6 MΩ was achieved. Unless otherwise stated, all voltage-clamp recordings were performed at a holding potential of -65 mV and all current-clamp recordings from the resting membrane potential (V_M_). For all recordings series resistance (R_S_) was monitored but not compensated in voltage-clamp and the bridge balanced following pipette-capacitance compensation in current-clamp. Signals were filtered online at 2-10 kHz using the built in 2-pole Bessel filter of the amplifier, digitized and acquired at 20 kHz (Digidata 1550B, Axon Instruments, USA), using pClamp 10 (Molecular Devices, CA, USA). Data was analysed offline using the open source Stimfit software package ((Guzman et al., 2014) http://www.stimfit.org). The liquid junction potential was measured as -12 mV and measurements not adjusted.

In whole-cell recordings the intrinsic properties of recorded neurons were characterized in current-clamp from resting membrane potential. A family of 500 ms hyper-to depolarising current steps (−250 to +250 pA, 50 pA steps; or -500 to +500 pA, 100 pA steps) were used depending on the initial input resistance of the neuron. Cells were identified on the basis of the voltage response and the resulting train of action potentials (AP) elicited by a family of hyper-to depolarizing current steps (50 pA, 500 ms duration; -500 pA to +500 pA). Neurons were rejected from further analysis if resting membrane potential was more depolarised than -50 mV, APs failed to overshoot 0 mV, initial access resistance (R_A_) exceeded 30 MΩ, or R_A_ fluctuated by >20% over the time course of the experiment.

### Characterization of postsynaptic GABA_B_R-mediated effects

To identify GABA_B_R mediated currents, neurons were recorded in the presence of ionotropic receptor blockers, CNQX or NBQX (10 µM), DL-AP5 (50 µM) and either picrotoxin (50 µM) or SR-95531 (Gabazine, 10 μM), which were bath applied. Extracellular stimuli were delivered via a bipolar twisted Ni:Chrome wire electrode placed at the border of L1 and L2. GABA_B_R-mediated IPSCs were evoked by 200 Hz trains of 5 stimuli (Booker *et al*., 2013; Booker *et al*., 2020) and a minimum of 15 IPSCs collected. The amplitude of GABA_B_R-mediated IPSCs was measured in average traces (>5 traces) over a 10 ms peak, within a 200 ms region relative to the pre-stimulus baseline. GABA_B_R-mediated whole-cell currents (I_WC_) were assessed by 5 to 10-minute bath application of the canonical agonist R-baclofen (10 µM). To confirm that baclofen-induced I_WC_ was GABA_B_R mediated, the potent and selective antagonist CGP 55,845 (CGP, 5 μM) was then applied to the bath. Peak baclofen and CGP currents were measured as the baseline I_WC_ over a 2-minute window of each pharmacological epoch, relative to baseline.

### Pharmacological manipulation of neuronal discharge

For baclofen puff experiments, a patch pipette was filled with 150 mM NaCl containing 50 μM baclofen and lowered to the slice adjacent to the recorded cell. Upon successful establishment of a whole-cell recording, in the presence NBQX (10 µM), DL-AP5 (50 µM) and either picrotoxin (50 µM) or SR-95531 (Gabazine, 10 μM), the puff pipette was placed just below the surface of the slice, approximately 50-100 μm distal from the recorded cell. Based on intrinsic characterisation of the cell (see above), the rheobase was identified and a +25 pA bias current applied for baclofen puffs. Each puff was delivered via a PicoSpritzer (10 mbar, 200 ms) at 1-minute intervals. In most cells, 1-3 baclofen puffs were applied under voltage-clamp (−65 mV) to determine the current amplitude evoked. Following successful measurement of AP discharge following baclofen puffs, CGP was applied to the bath to confirm that inhibition was baclofen mediated. For analysis, spontaneous APs detected and frequency calculated in 200 ms temporal bins, plotted against time. The degree of perisomatic inhibition was calculated 2-3 s after the onset of puff application.

### Characterization of presynaptic GABA_B_R-mediated effects

Monosynaptic excitatory postsynaptic currents (EPSC) were examined in human and rat L2/3 and L5 neurons in the presence of picrotoxin (50 µM) added to the perfusing ACSF. To evoke synaptic responses, extracellular stimuli were delivered via a Ni:Chrome twisted bipolar electrode placed in either L1 and L4 (for L2/3 PCs) or L1 and L5 (for L5 PCs). EPSCs were elicited using paired stimulus (2x stimuli at 20 Hz) repeated at 0.1 Hz. Stimulus intensity was titrated to produce a monosynaptic response of ∼100 pA. After 5 minutes of stable recording, the GABA_B_R agonist baclofen was applied to the bath at 10 µM. Following steady-state of baclofen wash-in, we then applied the potent and selective GABA_B_R antagonist CGP-55,845 (5 µM) to confirm receptor specificity. The amplitude of EPSCs was measured over a 10 ms window following the stimulus artifact and mean data are presented as the average of 10 traces normalised to baseline levels over the 2 min prior to baclofen wash-in. For PPR comparisons, the second EPSC amplitude was divided by the first.

### In vitro oscillations

For local field potential (LFP) recordings, slices were transferred to an interface recording chamber which perfused with carbogenated recording ACSF at 2-3 ml/min, maintained at 30 ± 1°C. Recording pipettes with a resistance of 1-3 MΩ were pulled from borosilicate glass capillaries (1.5 mm outer/0.86 mm inner diameter, Harvard Apparatus, UK) on a horizontal electrode puller and filled with recording ACSF. Slices were visualized by means of a wide-field microscope (Leica, Germany) and pipettes placed in L2/3 and L5. To induce population level oscillations, kainate (KA, 0.6 - 2 µM) and carbachol (CCh, 60-200 µM) were applied to perfusing ACSF for a minimum of 30 minutes. Baclofen (2 and 20 μM) or CGP (5 μM) were applied to the perfusing ACSF, additional to KA and CCh. LFP recordings were accomplished using a 2-channel amplifier (EXT-02 B, NPI Electronics, Germany). Signals were low-pass (1 Hz) and high-pass (500 Hz) filtered online, and acquired at 20 kHz (Micro1401-3, Cambridge Electronic Design, UK). All raw data were collected using Spike2 (Cambridge Electronic Design, UK). For analysis, 2-minute epochs were notch filtered at (49-51 Hz), bandpass filtered from 1-100 Hz, then fast-Fourier transform analysis performed to generate power spectra. Signals were rejected from analysis if epileptiform-like activity was present following induction of oscillatory activity or if beta/gamma (12-100 Hz) power was lower than 0.4 µV^2^.

### LFP / AUC

Fast-Fourier transforms (FFTs) were performed on 2-minute epochs of filtered signal to determine the amplitude (µV^2^) and frequency (Hz) of each signal following 30-40 minutes KA+CCh, 2µM and 20µM baclofen application. Employing a resolution of 0.61 Hz within a Hanning window, FFT size generated 164 data points spanning 0.61-99.4 Hz, capturing power and corresponding frequency. FFT amplitude values were normalised to the frequency resolution (0.61 Hz) yielding a power spectral density plot illustrating amplitude as a function of frequency (µV²/Hz). Frequency bands were defined as theta (4-10 Hz), beta (10-30 Hz), low-gamma (30-50 Hz), and high-gamma (50-85 Hz). Area under the curve (AUC) was calculated as the amplitude of each frequency band between lower and upper bandwidths during control (KA+CCh) or baclofen conditions. AUC values were normalized to KA+CCh to quantify amplitude changes post-baclofen application.

### Auto- and cross-correlation analysis

The same 2-minute epochs used in spectral analysis were imported into MATLAB and analysed using custom MATLAB scripts. Signals were subdivided into three 10-second epochs (0-10, 50-60, and 110-120 seconds) then bandstop filtered across the four distinct frequency bands. Auto-correlation used the Xcorr() function to determine the correlation strength of each frequency band over time by comparing a static signal to a time-lagged copy of itself. Cross-correlation similarly used the Xcorr() function to gauge the strength between prominent frequency pairs following KA+CCh and baclofen conditions in all L2/3 and L5 recordings. Cross-correlation between distinct frequency bands were measured from slices with simultaneously oscillating L2/3 and L5 recordings. Representative correlograms were generated to produce a visualisation of the coupling between oscillations. The strength of cross-correlation analysis was quantified using the peak value at zero-time lags.

### Phase-amplitude coupling analysis

The strength of phase-amplitude coupling (PAC) between distinct frequency bands was measured using a custom MATLAB script, based on earlier studies (Tort *et al*., 2010). 2-minute epochs of 50 Hz bandpass filtered signal were imported into MATLAB. Signals were zero-phase filtered using the filtfilt() function, then bandstop filtered to isolate each distinct frequency band. Hilbert transforms were run to generate a normalised phase-amplitude plot of the isolated lower frequency phase angles and the amplitude envelope of the higher frequency oscillation. The modulation index measure was used to quantify the PAC strength, calculating the deviations of phase-amplitude from a uniform distribution, with a smaller variation producing a larger modulation index value indicative of stronger PAC.

### Visualization, imaging and reconstruction of the recorded neurons

Recorded neurons were visualised as previously described (Booker et al., 2014). Following successful experiments, recorded cells were sealed with outside-out patch formation and then fixed in 4 % paraformaldehyde (PFA) diluted in 0.1 M phosphate buffer (PB) for 24-48 hours at 4°C. Slices were rinsed in PB, then phosphate buffered saline (PBS; 0.1 M PB + 0.9 % NaCl). Slices were then incubated for 48-72 hours in a PBS solution containing 0.3-0.5% TritonX-100 and 0.05% NaN_3_ and streptavidin conjugated to either AlexaFluor568 or AlexaFluor633 (1:500; Invitrogen, UK). Slices were then washed in PBS, then PB, and mounted on glass slides with a hard-set mounting medium (Vectashield, Vector Labs, UK) and then cover-slipped. Biocytin filled cells were imaged with a laser scanning confocal microscope (SP5, Leica, Germany) using a 20x objective and z-axis stacks of images collected (2048×2048 pixel radial resolution, 1 μm axial steps). Example cells were reconstructed off-line from image stacks digitally stitched and segmented using semi-automatic analysis software (Simple Neurite Tracer plug-in for the FIJI software package (http://fiji.org); (Longair et al., 2011)

### Statistical Analysis

Statistical comparisons were made using either Mann-Whitney or Wilcoxon matched-pairs tests, 1-, 2-, or 3-way ANOVAs, and linear regression (Pearson’s tests). ANOVAs were further analysed using Holm-Sidak post-tests, adjusted for multiple comparisons. All statistical analysis was performed using GraphPad Prism. For group-wise comparisons data is shown as box-plots with median and 25-75% percentiles displayed in the box; minimum and maximum extents are shown as whiskers. For most rats and human cases 1-2 cells per cell type and experiment were collected. As such, it was not possible to accurately calculate intra-replicate variability. As such, all data is shown and analysed as cell averages. For current-frequency, time-course, and spectral plots, the mean ± SEM are displayed. All tables state the mean ± SD of observed data. Statistical significance was assumed if P<0.05. A full summary of all reported values and statistical analysis is provided in the supplementary materials.

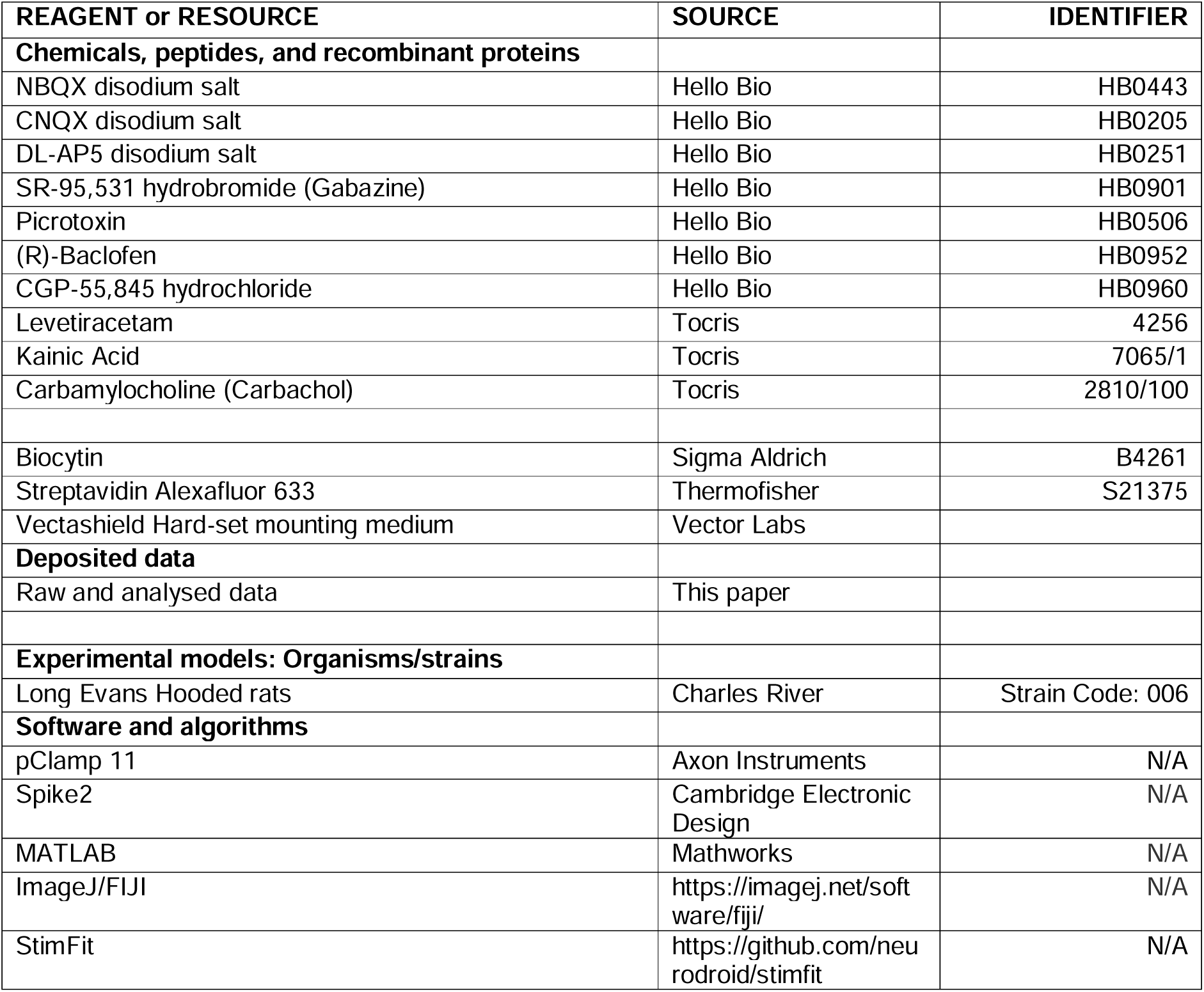
Key resources table

| REAGENT or RESOURCE | SOURCE | IDENTIFIER |
| --- | --- | --- |
| **Chemicals, peptides, and recombinant proteins** |  |  |
| NBQX disodium salt | Hello Bio | HB0443 |
| CNQX disodium salt | Hello Bio | HB0205 |
| DL-AP5 disodium salt | Hello Bio | HB0251 |
| SR-95,531 hydrobromide (Gabazine) | Hello Bio | HB0901 |
| Picrotoxin | Hello Bio | HB0506 |
| (R)-Baclofen | Hello Bio | HB0952 |
| CGP-55,845 hydrochloride | Hello Bio | HB0960 |
| Levetiracetam | Tocris | 4256 |
| Kainic Acid | Tocris | 7065/1 |
| Carbamylocholine (Carbachol) | Tocris | 2810/100 |
| Biocytin | Sigma Aldrich | B4261 |
| Streptavidin Alexafluor 633 | Thermofisher | S21375 |
| Vectashield Hard-set mounting medium | Vector Labs |  |
| **Deposited data** |  |  |
| Raw and analysed data | This paper |  |
| **Experimental models: Organisms/strains** |  |  |
| Long Evans Hooded rats | Charles River | Strain Code: 006 |
| **Software and algorithms** |  |  |
| pClamp 11 | Axon Instruments | N/A |
| Spike2 | Cambridge Electronic Design | N/A |
| MATLAB | Mathworks | N/A |
| ImageJ/FIJI | https://imagej.net/software/fiji/ | N/A |
| StimFit | https://github.com/neurodroid/stimfit | N/A |

## Supplemental information titles and legends

*Supplementary Table 1: Summary of human case details.* A summary of the key clinical features of the patients consented for the current study, according to age. Listing the sex at birth, brain region, hemisphere, reason for surgery, history of seizures, prescription of Levetiracetam (LEV), and number of replicates included in this study from whole-cell (WC) and local field potential (LFP) recordings.

*Supplementary Table 2: Electrophysiological properties of rat and human neurons*. Key measurements made from all recorded neurons in 1-month (1 m), 6+ month (6+ m), and adult humans from L2/3 and L5. Data is shown as mean ± SD. Statistics shown as p-values from Student’s T-tests, before and after multiple comparison adjustment.

*Supplementary Figure 1: Electrophysiological properties of human L2/3 neurons minimally correlates with age.* Correlation analysis of age (years) and key electrophysiological properties of identified human L2/3 neurons. Note few cellular physiological properties closely correlated with age. Other key parameters show strong correlations in clusters, in particular passive membrane and AP properties.

*Supplementary Figure 2: Presynaptic GABA_B_R-mediated inhibition of synaptic inputs to 1 month old rats.* (**A**) Example EPSCs evoked by L1 stimulation in L2/3 PCs recorded at -70 mV voltage clamp under control conditions (Ctrl) and following bath application of 10 μM baclofen (Bacl.) and 5 μM CGP-55,845 (CGP). Control recordings are shown for reference (grey traces). Lower, time-course of EPSC amplitude following baclofen (solid bar) and CGP (dashed bar) wash-in. Control baseline is shown for reference (grey dashed line). (**B**) Data in the same form but for L1 inputs to L5 PCs (blue). (**C**) Data in the same form, but for L4 inputs to L2/3 PCs (light grey). (**D**) Data in the same form, but for L5 inputs to L5 PCs (light blue). Data is shown as mean ± SEM.

*Supplementary Figure 3: Presynaptic GABA_B_R-mediated inhibition of synaptic inputs to 6+ month old rats*. (**A**) Example EPSCs evoked by L1 stimulation in L2/3 PCs recorded at -70 mV voltage clamp under control conditions (Ctrl) and following bath application of 10 μM baclofen (Bacl.) and 5 μM CGP-55,845 (CGP). Control recordings are shown for reference (grey traces). Lower, time-course of EPSC amplitude following baclofen (solid bar) and CGP (dashed bar) wash-in. Control baseline is shown for reference (grey dashed line). (**B**) Data in the same form but for L1 inputs to L5 PCs (blue). (**C**) Data in the same form, but for L4 inputs to L2/3 PCs (light grey)**. (D**) Data in the same form, but for L5 inputs to L5 PCs (light blue). Data is shown as mean ± SEM.

*Supplementary Figure 4: Presynaptic short-term plasticity is variably modified by baclofen at rat cortical glutamatergic synapses, but is robustly controlled in human cortex.* (**A**) Quantification of PPR before (Ctrl) and after bath application of 10 μM baclofen (Bacl.) or 5 μM CGP-55,845 (CGP) at L1 (black) and L4 (grey) inputs to L2/3 PCs (n=8 cells) and L1 (blue) and L5 (light blue) inputs to L5 PCs (n=7 cells) in 1-month old rats. (**B**) Data in the same form for 6+ month old rats (n=6 L2/3 cells, n=5 L5 cells). (**C**) Data in the same form for 28-77 year old human cortex (n=9 L2/3 cells, n=9 L5 cells). All data is shown as mean ± SEM (due to pairwise nature) and is shown with results from individual cells. Statistics shown: ns – p>0.05, * - *p*<0.05, ** - *p*<0.01 *** - *p*<0.001, all from Holm-Sidak post-tests.

*Supplementary Figure 5: Comparison of paired-pulse ratios at synaptic inputs to cortical neurons reveals greater modulation in human cortex.* Paired pulse ratio (PPR) of EPSCs at L1 (**A**, black) and L4 (**B**, grey) inputs to L2/3 PCs or L1 (**C**, blue) and L5 (**D**, light blue) inputs to L5 PCs; in 1- and 6+ months old rats, and humans. (**E**) Comparison of baclofen-mediated modulation of PPR at synaptic inputs to L2/3 PCs expressed as % of control (ctrl) reveals greater modulation of human synapses either from L1 (black) or L4 (grey). (**F**) Comparable modulation of EPSC PPR at L1 (blue) or L5 (light blue) inputs to L5 PCs, regardless of species. All data is shown as box plots, with data from individual cells shown overlaid (open circles). Statistics shown from 1-way ANOVA with Holm-Sidak post-tests (A, B, C, D) or 2-way ANOVA (E, F); ns – p>0.05, * - *p*<0.05.

*Supplementary Figure 6. Human L5 displays reduced correlation following baclofen bath application.* Example plots of cross-correlation strength between prominent oscillations in LFP recordings from human L5 under control conditions (blue) and following 2 µM (light blue) or 20 µM (lightest blue) baclofen bath application.

*Supplementary Figure 7: Increased sensitivity of human neuronal oscillations to baclofen, as compared to 6+ month old rats*. (**A**) Subtracted power spectra from fast-Fourier transforms of filtered (1-100 Hz bandpass) LFP recordings from L2/3 of human neocortex (black) and 6+ month rat S1 (green), both under control conditions (KA+CCh, black). (**B**) Power spectra of L2/3 human and S1 6+ month rat cortex following 2 µM (grey, green) or 20 µM (light grey, light green) baclofen (Bacl.) bath application, respectively. (**C**) Integrals of each frequency epoch, shown as area-under-curve (AUC), for human (black) and rat (green) control LFP recordings revealed no difference in overall power (F_(1, 44)_ = 0.64, *p*=0.427, 2-way ANOVA [species]). (**D**) Comparison of the effect of baclofen on oscillatory power for human (black) and rat (green) LFP recordings. Data is shown as the % change from control AUC for each frequency band, with respect to either 2 μM (left) and 20 μM (right) baclofen, with a statistically robust effect of species on the degree of inhibition (F_(1, 72)_ = 11.4, *p*=0.0012 [species], 3-way ANOVA). Data are shown as mean ± SEM (A, B), or box-plots (min-max; C, D) with individual data point overlain.

*Supplementary Figure 8: Cross-correlation strength of prominent oscillations between cortical layers reveals minimal effect of baclofen in vitro.* (**A**) Example cross-correlograms showing the strength of interaction between similar frequency oscillations under control conditions (KA+CCh, Black) and following 2 µM (grey) and 20 µM (light grey) baclofen application, with respect to lag time. (**B**) Grouped cross-correlation data showing positive and negative correlation across simultaneous oscillatory activity under control conditions, and following 2 µM (grey) and 20 µM (light grey) baclofen application revealed no effect of baclofen (F_(2, 66)_ = 0.26, *p*=0.77, 2-way ANOVA). Data are shown as mean ± SEM with values from individual slices overlaid.

*Supplementary Figure 9: Phase amplitude coupling in adult human brain slices reveals oscillatory coupling persists in the presence of baclofen.* (**A**) Example phase-amplitude plots of 2 minute signal epochs of L2/3 under control (KA+CCh, left) and following 2 µM (middle) and 20 µM (right) baclofen application. Phase amplitude is shown as colour coded as low (blue) to high (red) amplitude coupling strength, and is unitless. (**B**) Similar representation as in **A**, but in L5 recordings. **C**) Example phase-amplitude plots reflecting the strength (unitless) between oscillations as a function of the phase of the lower frequency oscillation between prominent oscillations in L2/3 control conditions (solid, black) and after 2 µM (dashed, light grey) and 20 µM (solid, dark grey) baclofen application. (**D**) The same as **C**, but for L5 recordings under control conditions (solid, blue) and after 2 µM (dashed, blue) and 20 µM (solid, light blue) **E**) Peak modulation index amplitudes between each frequency band in L2/3 under control (black) and following 2 µM (grey) and 20µM (light grey) baclofen application. (**F**) The same data, but for L5. Data is shown as mean ± SEM (E, F).

