## Supplementary Materials for "Phylogenetic divergence of GABA_B_ receptor signalling in neocortical networks over adult life"

| Age | Sex | Diagnosis | Hemisphere | Region | Seizures? | LEV | WC | LFP |
| --- | --- | --- | --- | --- | --- | --- | --- | --- |
| 28 | F | Glioblastoma | Left | Temporal | Yes | Yes | 3 | 1 |
| 30 | M | Glioblastoma | Left | Frontal | No | No | 2 |  |
| 32 | M | Glioma | Left | Frontal | No | No | 3 | 1 |
| 32 | F | Astrocytoma | Left | Temporal | Yes | Yes | 4 |  |
| 32 | F | Glioma | Left | Frontal | Yes | Yes | 3 |  |
| 37 | F | Glioma | Right | Frontal | No | No | 3 |  |
| 39 | F | Glioma | Left | Frontal | Yes | Yes | 1 |  |
| 39 | F | Breast cancer with metastases | Left | Frontal | Yes | Yes | 4 | 1 |
| 43 | F | Probable Glioblastoma | Right | Frontal | No | No | 2 | 1 |
| 45 | M | Glioblastoma | Right | Frontal | Yes | Yes |  |  |
| 47 | M | Glioma | Right | Parietal | Yes | Yes | 4 | 1 |
| 48 | F | Glioma | Left | Frontal | No | Yes | 3 | 1 |
| 50 | M | Gliosarcoma | Left | Temporal | No | No | 1 |  |
| 50 | M | Glioma | Left | Frontal | No | No | 7 |  |
| 50 | M | Glioma | Right | Frontal | Yes | Yes | 6 |  |
| 51 | M | Glioblastoma | Right | Frontal | No | No |  | 1 |
| 52 | F | Metastases | Left | Frontal | Yes | Yes |  | 1 |
| 53 | M | Renal metastases | Right | Temporal | No | No | 2 |  |
| 54 | M | Glioma | Left | Frontal | No | Yes |  | 1 |
| 55 | M | Glioblastoma | Right | Temporal | Yes | Yes | 17 |  |
| 55 | M | Glioblastoma | Right | Frontal | No | Yes | 3 | 1 |
| 56 | M | Glioma | Left | Parietal | Yes | Yes | 3 |  |
| 59 | M | Glioblastoma | Right | Parietal | No | No |  | 1 |
| 60 | F | Glioblastoma | Right | Frontal | No | No | 3 |  |
| 60 | F | Glioma | Left | Occipital | Yes | Yes | 4 |  |
| 60 | F | Glioma | Left | Parietal | Yes | Yes | 2 |  |
| 60 | M | Lymphoma | Left | Frontal | Yes | Yes | 7 |  |
| 63 | M | Glioblastoma | Left | Temporal | Yes | Yes | 2 |  |
| 63 | F | Lung cancer with metastases | Left | Frontal | No | No |  | 1 |
| 64 | M | Malignant melanoma | Right | Temporal | No | No | 2 |  |
| 64 | M | Lung cancer with metastases | Left | Frontal | No | No |  | 1 |
| 65 | M | Glioblastoma | Right | Frontal | No | No | 1 |  |
| 65 | M | Glioblastoma | Left | Frontal | No | No | 5 |  |
| 65 | M | Glioma | Left | Temporal | No | No | 7 | 1 |
| 65 | M | Glioma or Metastases | Right | Frontal | No | No | 3 |  |
| 65 | M | Glioblastoma | Left | Temporal | No | Yes | 3 |  |
| 66 | M | Glioblastoma | Left | Parietal | Yes | Yes | 3 | 1 |
| 66 | F | Glioma | Right | Parietal | Yes | Yes | 1 |  |
| 68 | F | Glioblastoma | Left | Temporal | No | No | 1 |  |
| 70 | M | Glioma | Bifrontal | Frontal | Yes | Yes |  | 1 |
| 72 | M | Glioblastoma | Left | Frontal | No | No | 3 |  |
| 72 | M | Probable Glioblastoma | Right | Frontal | No | No | 2 |  |
| 72 | M | Glioblastoma | Right | Parietal | No | No | 3 | 1 |
| 74 | F | Glioblastoma | Right | Temporal | No | No | 3 |  |
| 74 | M | Glioblastoma | Right | Temporal | No | No | 4 |  |
| 75 | M | Glioblastoma | Left | Frontal | No | No | 2 |  |
| 76 | F | Intracranial metastases | Right | Temporal | Yes | Yes | 4 | 1 |
| 77 | m | Glioblastoma | Right | Parietal | No | No |  | 1 |
| 77 | m | Glioblastoma | Left | Temporal | No | No | 4 |  |

*Supplementary Table 1: Summary of human case details.* A summary of the key clinical features of the patients consented for the current study, according to age. Listing the sex at birth, brain region, hemisphere, reason for surgery, history of seizures, prescription of Levetiracetam (LEV), and number of replicates included in this study from whole-cell (WC) and local field potential (LFP) recordings.

| Electrophysiological  Property | Human L2/3  n=33 cells  N=17 cases | Human L5  n=14 cells  N=8 cases | t | P | Adj. P |
| --- | --- | --- | --- | --- | --- |
| Membrane potential (mV) | -68.9 ± 5.8 | -68.1 ± 5.9 | 0.403 | 0.689 | 0.997 |
| Input resistance (MΩ) | 128.0 ± 104.9 | 111.8 ± 58.1 | 0.543 | 0.590 | 0.995 |
| Membrane time-constant (ms) | 16.6 ± 8.2 | 17.3 ± 8.5 | 0.269 | 0.790 | 0.997 |
| Capacitance (pF) | 175.7 ± 85.1 | 182.2 ± 91.5 | 0.231 | 0.818 | 0.997 |
| Rheobase (pA) | 194.7 ± 135.8 | 164.3 ± 84.2 | 0.775 | 0.443 | 0.995 |
| Voltage threshold (mV) | -40.3 ± 5.7 | -41.0 ± 7.6 | 0.346 | 0.731 | 0.997 |
| AP amplitude (mV) | 111.5 ± 10.8 | 104.4 ± 13.1 | 1.929 | 0.060 | 0.494 |
| AP 20-80% rise-time (ms) | 0.18 ± 0.06 | 0.19 ± 0.07 | 0.725 | 0.472 | 0.995 |
| AP half-height width (ms) | 0.82 ± 0.40 | 0.82 ± 0.36 | 0.069 | 0.945 | 0.997 |
| AP maximum decay (mV.ms^-1^) | 379.3 ± 126.7 | 326.3 ± 114.0 | 1.349 | 0.184 | 0.869 |
| AP maximum rise (mV.ms^-1^) | 110.9 ± 44.5 | 101.2 ± 35.4 | 0.726 | 0.472 | 0.995 |
| IF Slope (AP. pA^-1^) | 0.032 ± 0.015 | 0.049 ± 0.025 | 2.473 | 0.019 | 0.206 |

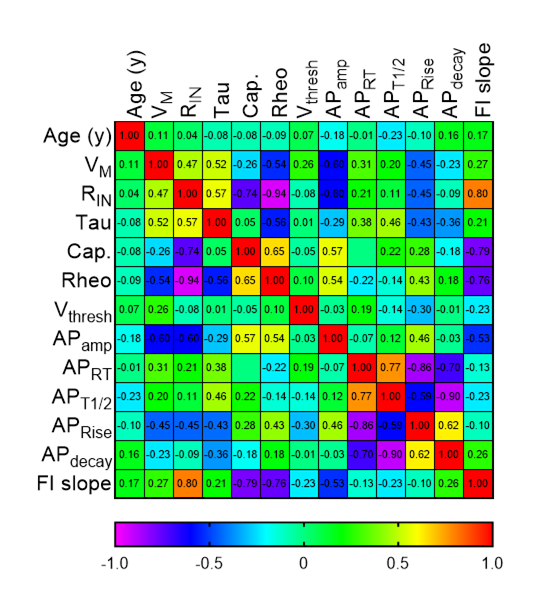
*Supplementary Table 2: Electrophysiological properties of rat and human neurons.* Key measurements made from all recorded neurons in 1 month (1 m), 6+ month (6+ m), and adult humans from L2/3 and L5. Data is shown as mean ± SD. Statistics shown as p-values from Student’s T-tests, before and after multiple comparison adjustment.

*Supplementary Figure 1: Electrophysiological properties of human L2/3 neurons minimally correlates with age.* Correlation analysis of age (years) and key electrophysiological properties of identified human L2/3 neurons. Note few cellular physiological properties closely correlated with age. Other key parameters show strong correlations in clusters, in particular passive membrane and AP properties.

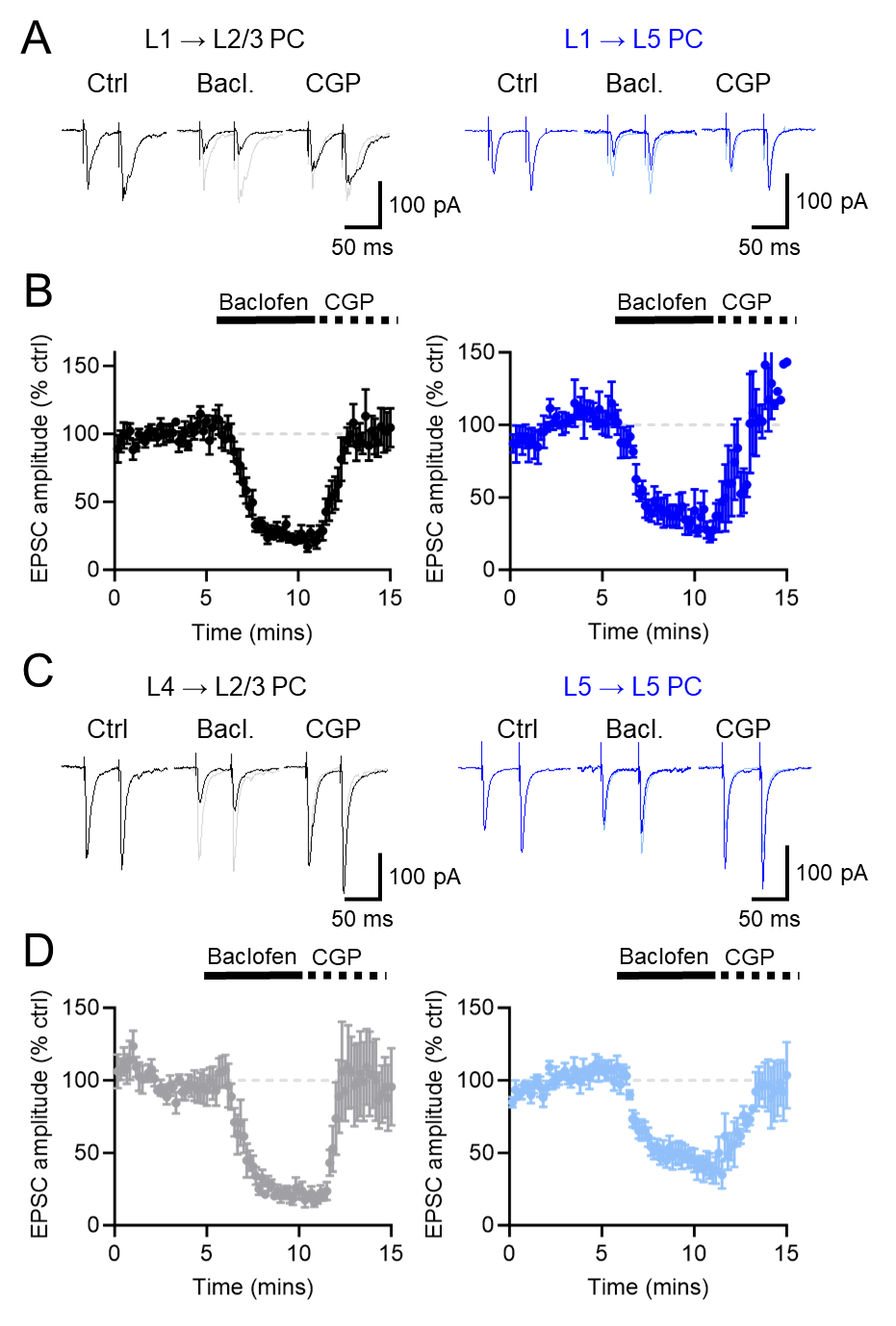
*Supplementary Figure 2: Presynaptic GABA_B_R-mediated inhibition of synaptic inputs to 1 month old rats*. **A)** Example EPSCs evoked by L1 stimulation in L2/3 PCs recorded at -70 mV voltage clamp under control conditions (Ctrl) and following bath application of 10 μM baclofen (Bacl.) and 5 μM CGP-55,845 (CGP). Control recordings are shown for reference (grey traces). Lower, time-course of EPSC amplitude following baclofen (solid bar) and CGP (dashed bar) wash-in. Control baseline is shown for reference (grey dashed line). **B**) Data in the same form but for L1 inputs to L5 PCs (blue). C) Data in the same form, but for L4 inputs to L2/3 PCs (light grey). **D**) Data in the same form, but for L5 inputs to L5 PCs (light blue). Data is shown as mean ± SEM.

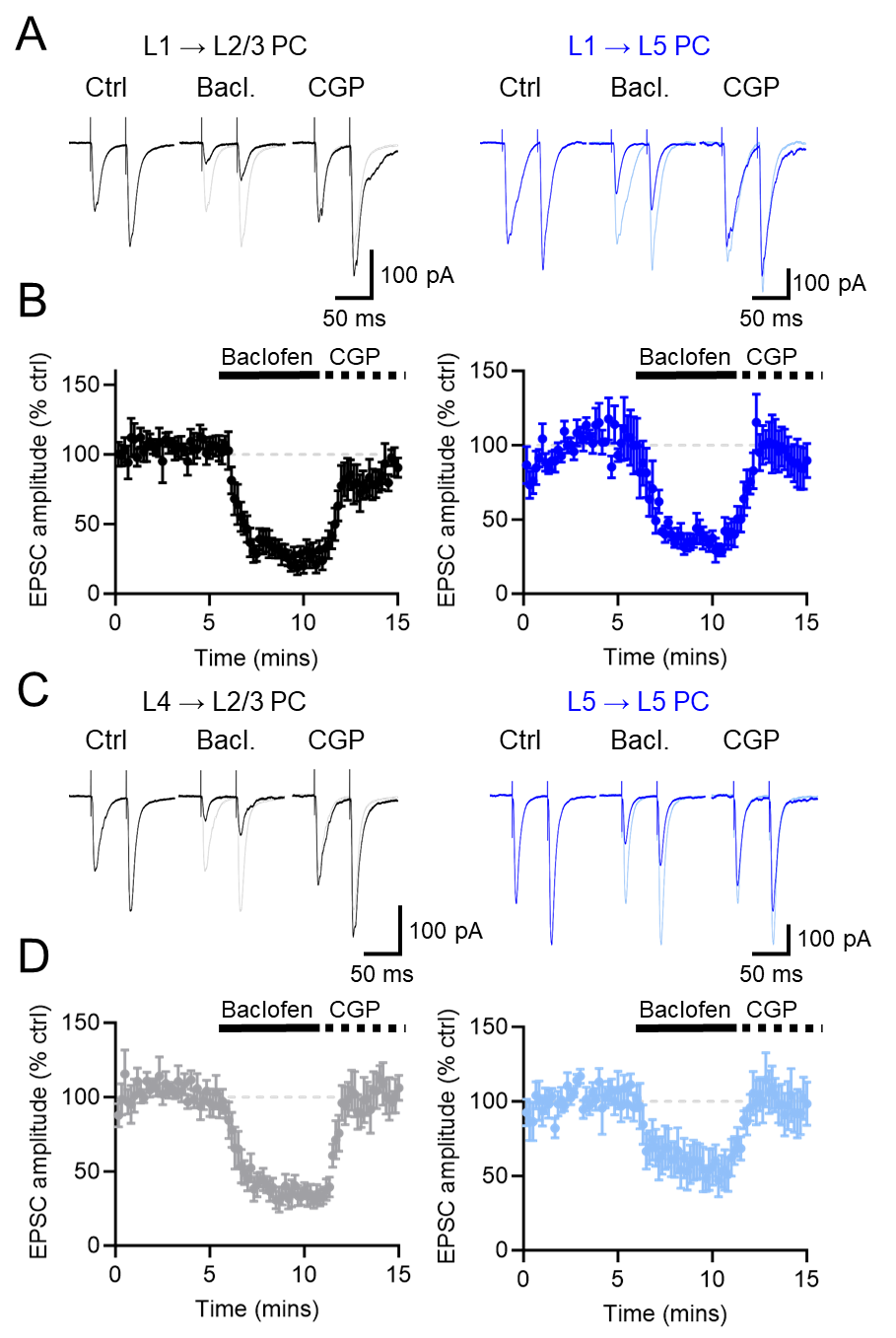
*Supplementary Figure 3: Presynaptic GABA_B_R-mediated inhibition of synaptic inputs to 6+ month old rats*. **(A**) Example EPSCs evoked by L1 stimulation in L2/3 PCs recorded at -70 mV voltage clamp under control conditions (Ctrl) and following bath application of 10 μM baclofen (Bacl.) and 5 μM CGP-55,845 (CGP). Control recordings are shown for reference (grey traces). Lower, time-course of EPSC amplitude following baclofen (solid bar) and CGP (dashed bar) wash-in. Control baseline is shown for reference (grey dashed line). (**B**) Data in the same form but for L1 inputs to L5 PCs (blue). (**C**) Data in the same form, but for L4 inputs to L2/3 PCs (light grey). (**D**) Data in the same form, but for L5 inputs to L5 PCs (light blue). Data is shown as mean ± SEM.

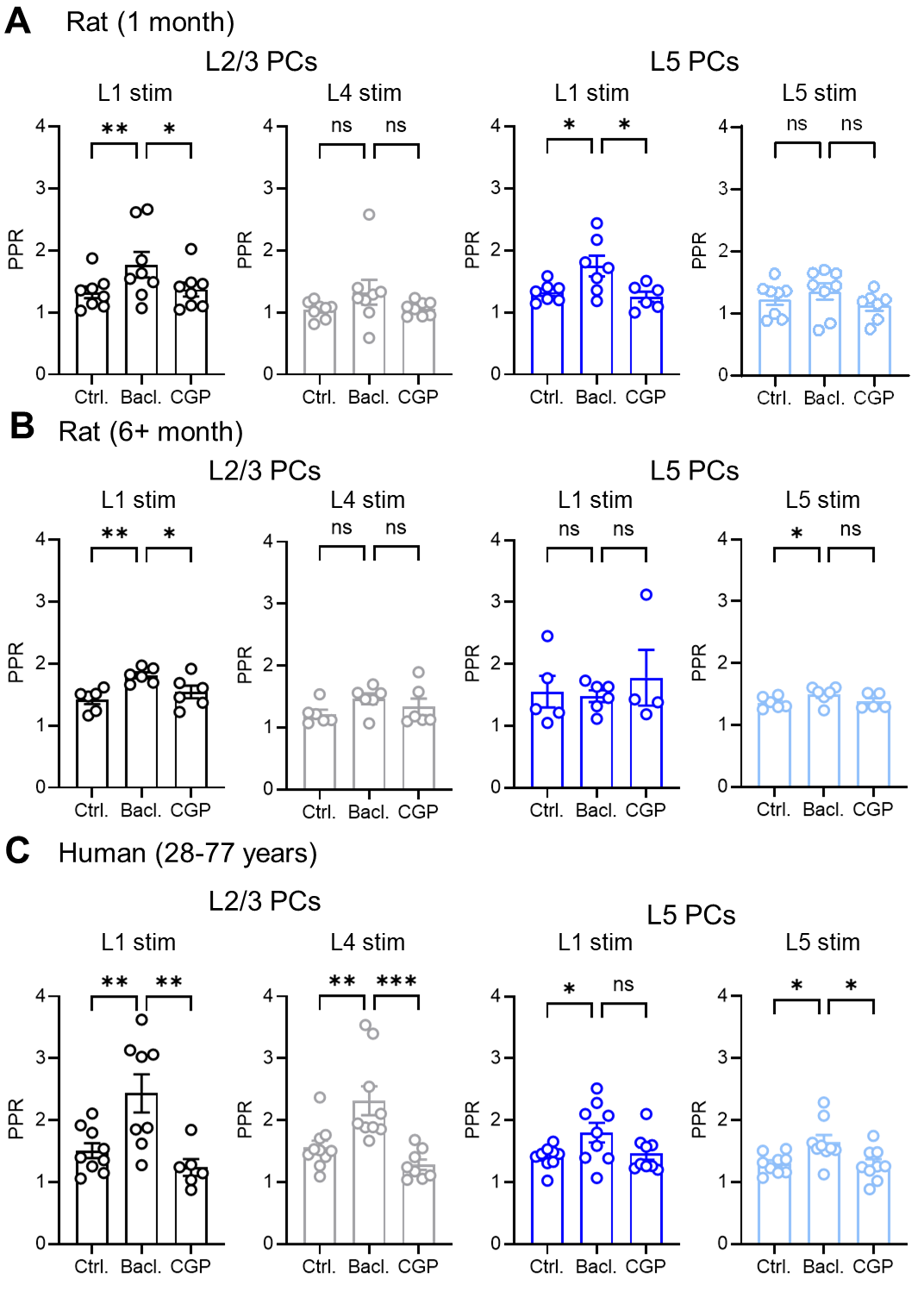

*Supplementary Figure 4: Presynaptic short-term plasticity is variably modified by baclofen at rat cortical glutamatergic synapses, but is robustly controlled in human cortex.* (**A**) Quantification of PPR before (Ctrl) and after bath application of 10 μM baclofen (Bacl.) or 5 μM CGP-55,845 (CGP) at L1 (black) and L4 (grey) inputs to L2/3 PCs (n=8 cells) and L1 (blue) and L5 (light blue) inputs to L5 PCs (n=7 cells) in 1 month old rats. (**B**) Data in the same form for 6+ month old rats (n=6 L2/3 cells, n=5 L5 cells). (**C**) Data in the same form for 28-77 year old human cortex (n=9 L2/3 cells, n=9 L5 cells). All data is shown as mean ± SEM (due to pairwise nature) and is shown with results from individual cells. Statistics shown: ns – p>0.05, * - *p*<0.05, ** - *p*<0.01 *** - *p*<0.001, all from Holm-Sidak post-tests.

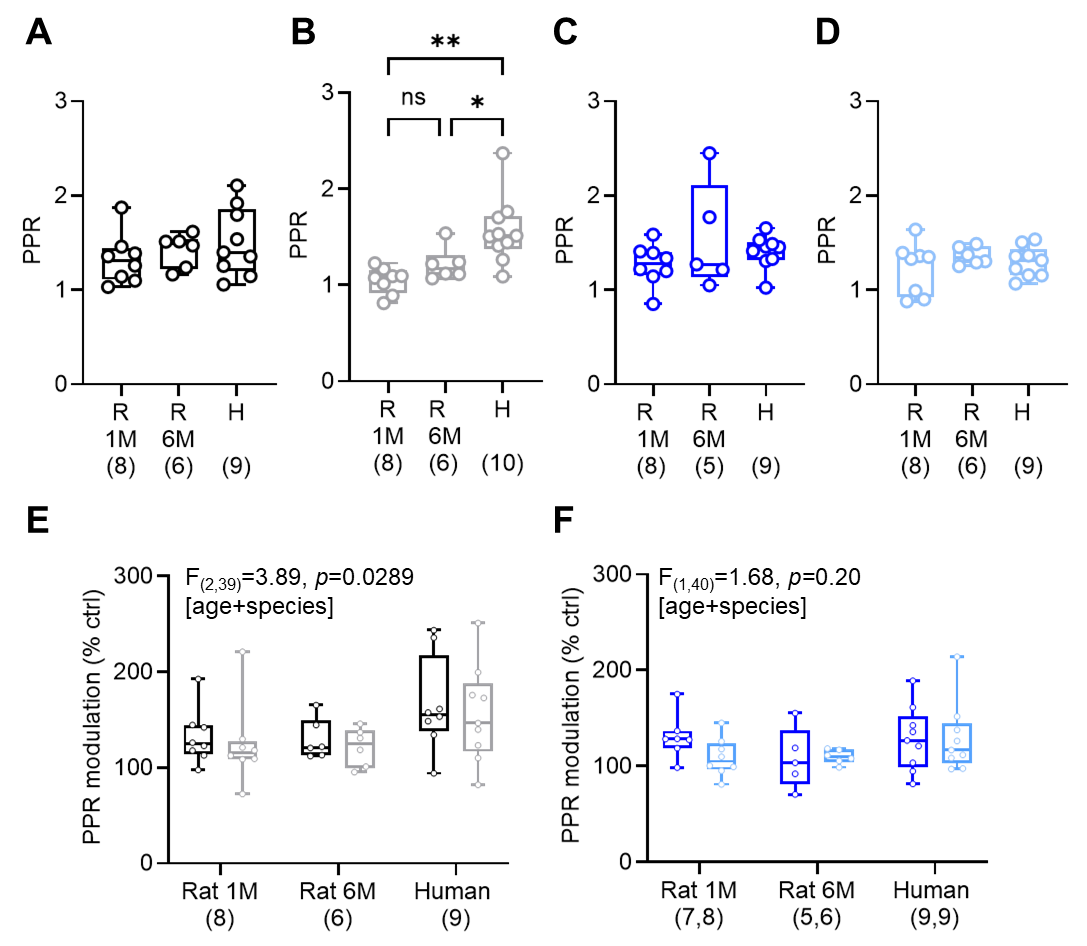
*Supplementary Figure 5: Comparison of paired-pulse ratios at synaptic inputs to cortical neurons reveals greater modulation in human cortex.* Paired pulse ratio (PPR) of EPSCs at L1 (**A**, black) and L4 (**B**, grey) inputs to L2/3 PCs or L1 (**C**, blue) and L5 (**D**, light blue) inputs to L5 PCs; in 1- and 6+ months old rats, and humans. (**E**) Comparison of baclofen-mediated modulation of PPR at synaptic inputs to L2/3 PCs expressed as % of control (ctrl) reveals greater modulation of human synapses either from L1 (black) or L4 (grey). (**F**) Comparable modulation of EPSC PPR at L1 (blue) or L5 (light blue) inputs to L5 PCs, regardless of species. All data is shown as box plots, with data from individual cells shown overlaid (open circles). Statistics shown from 1-way ANOVA with Holm-Sidak post-tests (A, B, C, D) or 2-way ANOVA (E, F); ns – p>0.05, * - *p*<0.05.

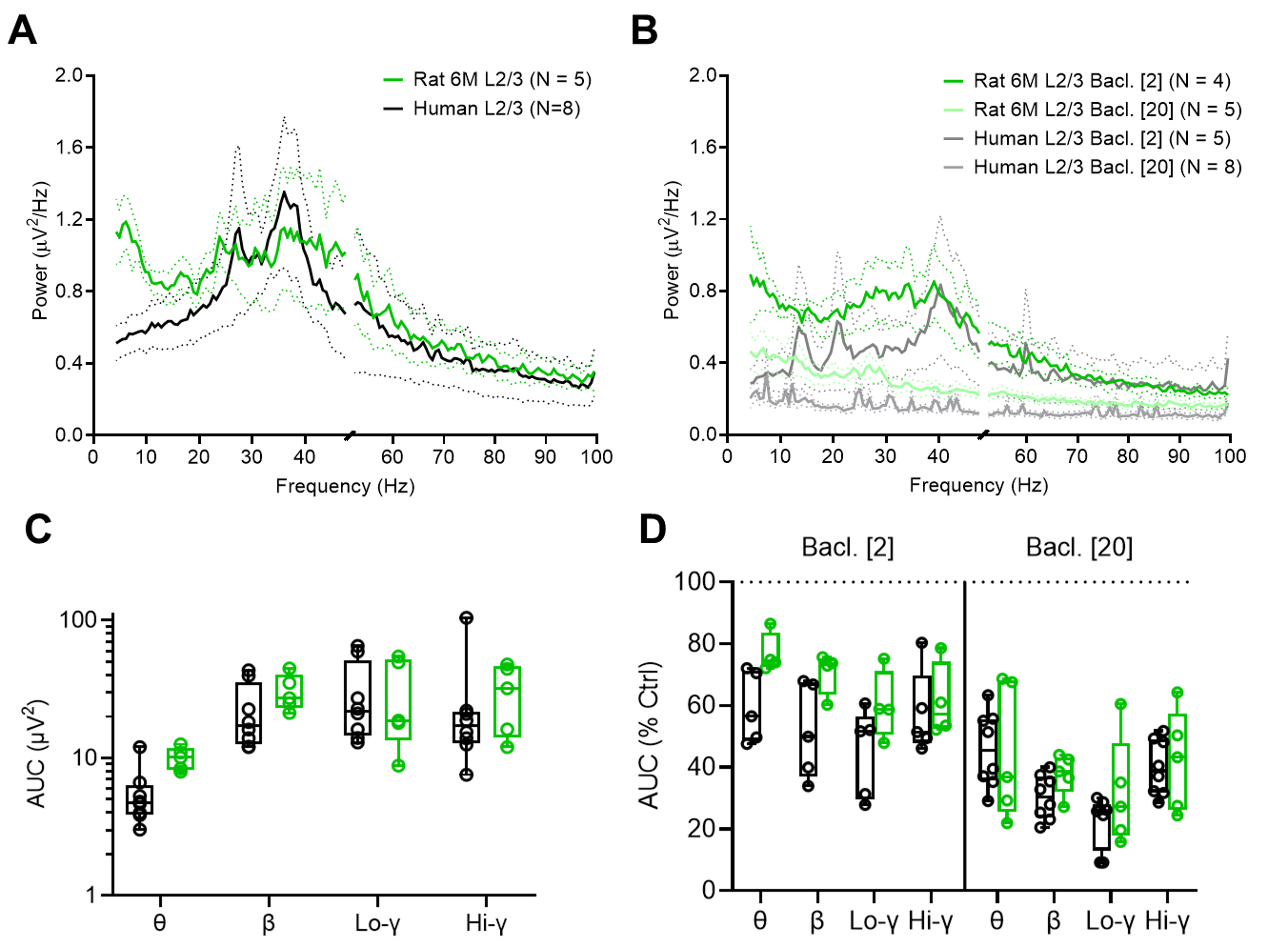

*Supplementary Figure 6: Increased sensitivity of human neuronal oscillations to baclofen, as compared to 6+ month old rats*. (**A**) Subtracted power spectra from fast-Fourier transforms of filtered (1-100 Hz bandpass) LFP recordings from L2/3 of human neocortex (black) and 6+ month rat S1 (green), both under control conditions (KA+CCh, black). (**B**) Power spectra of L2/3 human and S1 6+ month rat cortex following 2 µM (grey, green) or 20 µM (light grey, light green) baclofen (Bacl.) bath application, respectively. (**C**) Integrals of each frequency epoch, shown as area-under-curve (AUC), for human (black) and rat (green) control LFP recordings revealed no difference in overall power (F_(1, 44)_ = 0.64, *p*=0.427, 2-way ANOVA [species]). (**D**) Comparison of the effect of baclofen on oscillatory power for human (black) and rat (green) LFP recordings. Data is shown as the % change from control AUC for each frequency band, with respect to either 2 μM (left) and 20 μM (right) baclofen, with a statistically robust effect of species on the degree of inhibition (F_(1, 72)_ = 11.4, *p*=0.0012 [species], 3-way ANOVA). Data are shown as mean ± SEM (A, B), or box-plots (min-max; C, D) with individual data point overlain.

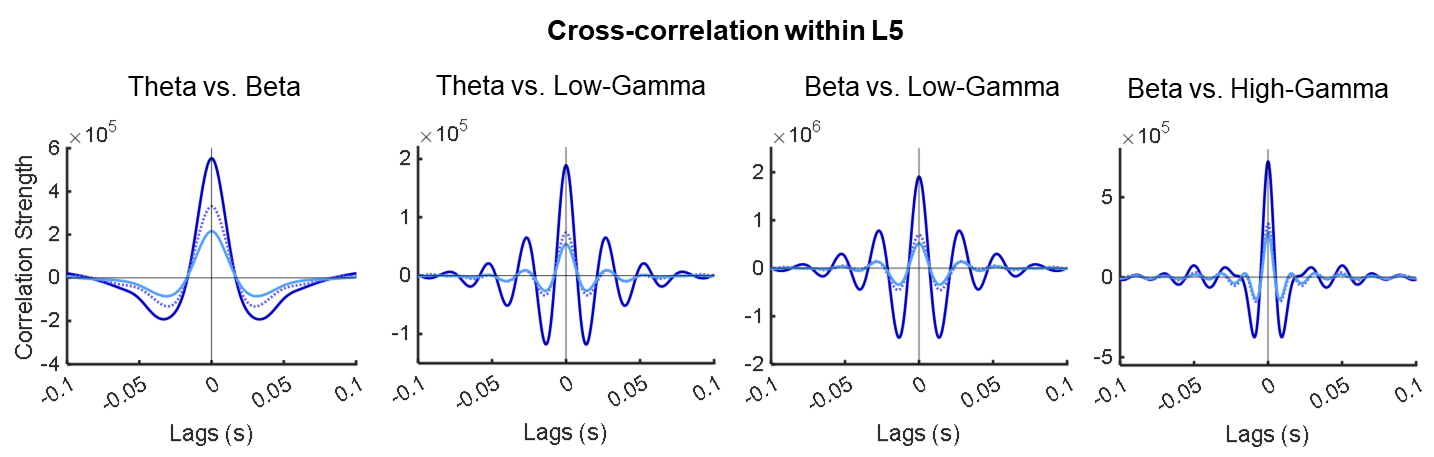
*Supplementary Figure 7: Human L5 displays reduced correlation following baclofen bath application.* Example plots of cross-correlation strength between prominent oscillations in LFP recordings from human L5 under control conditions (blue) and following 2 µM (light blue) or 20 µM (lightest blue) baclofen bath application.

*Supplementary Figure 8:* *Cross-correlation strength of prominent oscillations between cortical layers reveals minimal effect of baclofen in vitro.* (**A**) Example cross-correlograms showing the strength of interaction between similar frequency oscillations under control conditions (KA+CCh, Black) and following 2 µM (grey) and 20 µM (light grey) baclofen application, with respect to lag time. (**B**) Grouped cross-correlation data showing positive and negative correlation across simultaneous oscillatory activity under control conditions, and following 2 µM (grey) and 20 µM (light grey) baclofen application revealed no effect of baclofen (F_(2, 66)_ =
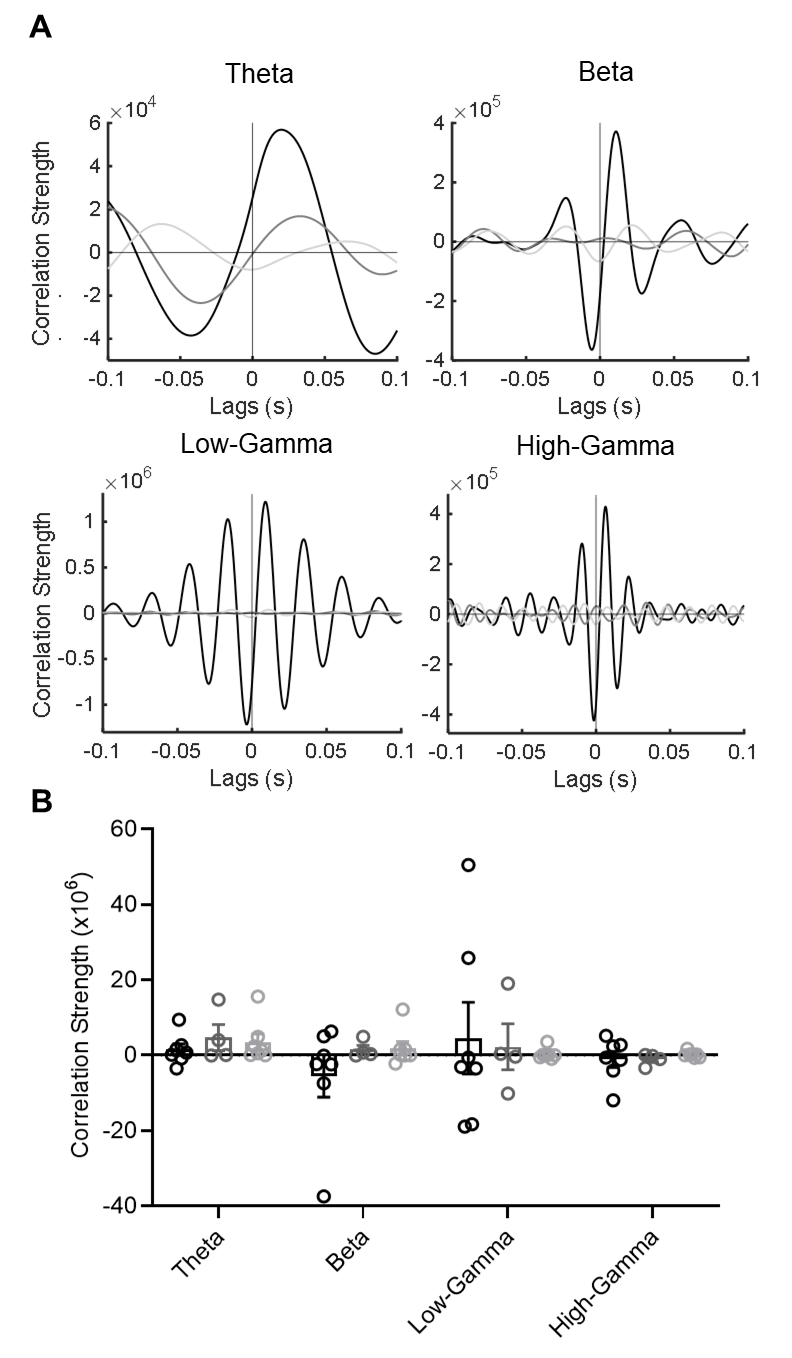
0.26, *p*=0.77, 2-way ANOVA). Data are shown as mean ± SEM with values from individual slices overlaid.

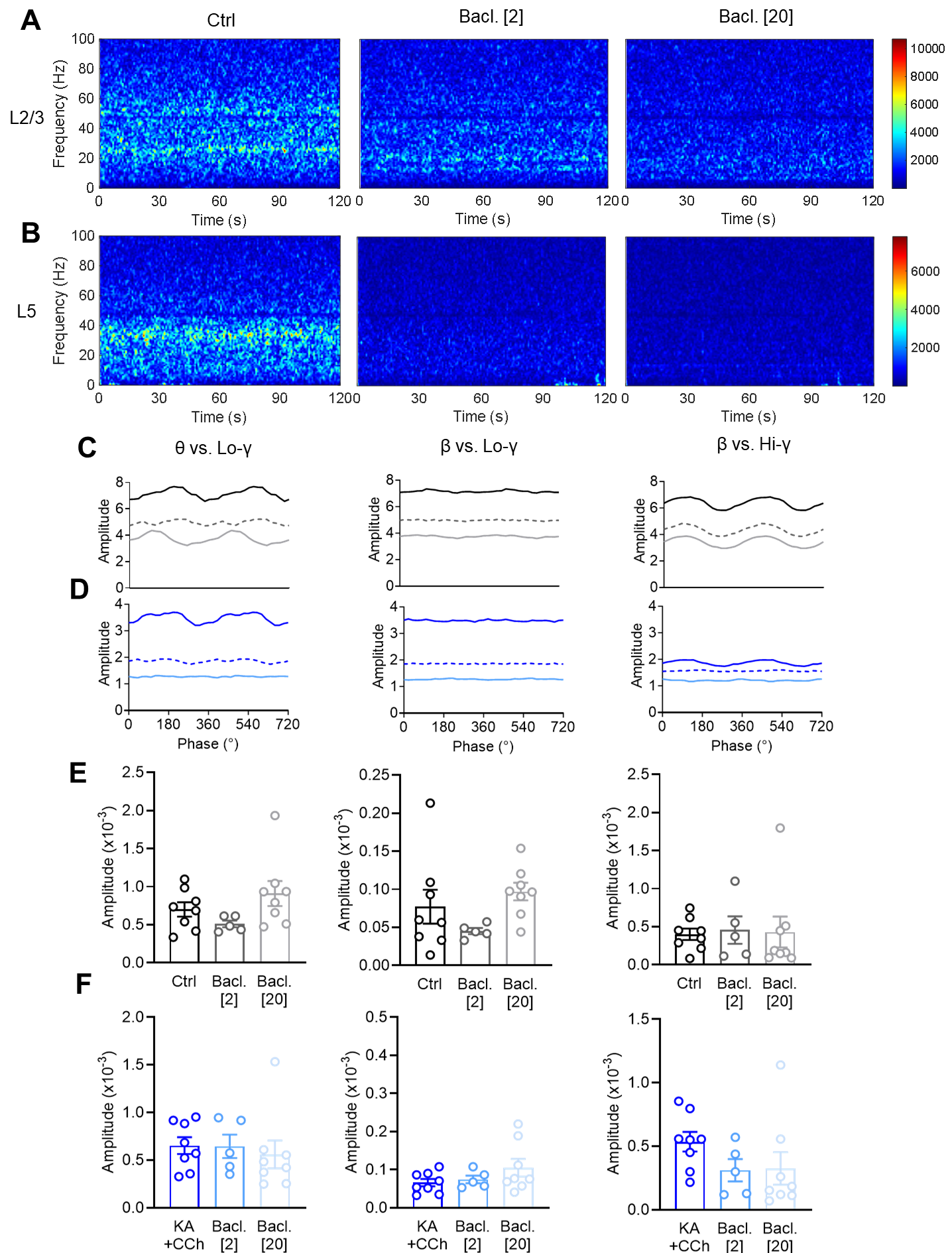

*Supplementary Figure 9: Phase amplitude coupling in adult human brain slices reveals oscillatory coupling persists in the presence of baclofen*. (**A**) Example phase-amplitude plots of 2 minute signal epochs of L2/3 under control (KA+CCh, left) and following 2 µM (middle) and 20 µM (right) baclofen application. Phase amplitude is shown as colour coded as low (blue) to high (red) amplitude coupling strength, and is unitless. (**B**) Similar representation as in **A**, but in L5 recordings. (**C**) Example phase-amplitude plots reflecting the strength (unitless) between oscillations as a function of the phase of the lower frequency oscillation between prominent oscillations in L2/3 control conditions (solid, black) and after 2 µM (dashed, grey) and 20 µM (solid, light grey) baclofen application. (**D**) The same as **C**, but for L5 recordings under control conditions (solid, blue) and after 2 µM (dashed, blue) and 20 µM (solid, light blue). (**E**) Peak modulation index amplitudes between each frequency band in L2/3 under control (black) and following 2 µM (grey) and 20µM (light grey) baclofen application. (**F**) The same data, but for L5. Data is shown as mean ± SEM (E, F).
